# Audiovisual integration is preserved in older adults across the cortical hierarchy

**DOI:** 10.1101/2023.01.28.526027

**Authors:** Samuel A. Jones, Uta Noppeney

**Affiliations:** Computational Neuroscience and Cognitive Robotics Centre, University of Birmingham, Birmingham, UK; Department of Psychology, Nottingham Trent University, Nottingham, UK; Donders Institute for Brain, Cognition & Behaviour, Radboud University, The Netherlands

## Abstract

Effective interactions with the environment rely on integration of multisensory signals: our brains must efficiently combine signals that share a common source, and segregate those that do not. Healthy ageing can change or impair this process. This functional magnetic resonance imaging study assessed the neural mechanisms underlying age differences in the integration of auditory and visual spatial cues. Participants were presented with synchronous audiovisual signals at various degrees of spatial disparity and indicated their perceived sound location. Behaviourally, older adults were able to maintain localisation accuracy, albeit with longer response times. At the neural level, they integrated auditory and visual cues into spatial representations along dorsal auditory and visual processing pathways similarly to their younger counterparts, but showed greater activations in a widespread system of frontal, temporal and parietal areas. According to multivariate Bayesian decoding, these areas encoded critical stimulus information beyond that which was encoded in the brain areas commonly activated by both groups. Surprisingly, however, the boost in information provided by these areas with age-related activation increases was comparable across the two age groups.

This dissociation—between comparable response accuracy and information encoded in brain activity patterns across the two age groups, but age-related increases in response times and regional activations—suggests that older participants accumulate noisier sensory evidence for longer, to maintain reliable neural encoding of stimulus-relevant information and thus preserve localisation accuracy.

## Introduction

The effective integration of multisensory signals is central to our ability to successfully interact with the world. Locating and swatting a mosquito, for example, relies on spatial information from hearing, vision, and touch. When signals from different senses are known to come from a common cause, humans typically perform this integration process in a statistically near-optimal way, weighting the contribution of each input by its relative reliability [1–5] (i.e. inverse of variance; though also see e.g. [6, 7]). However, determining specifically which signals share a common cause, and should thus be integrated, is computationally challenging. Young, healthy adults arbitrate between sensory integration and segregation in line with the predictions of normative Bayesian Causal Inference (BCI) [8–12]: they bind signals that are close together in space and time, but process signals independently when they are spatially or temporally disparate and hence unlikely to share a common source. Recent fMRI and EEG research has revealed that, for audiovisual spatial signals, these operations take place dynamically across the cortical hierarchy that encompasses primary sensory areas as well as higher-level regions such as intraparietal sulcus and planum temporale [10, 13]. Evidence also suggests that they interact with top-down attentional processes [5,14–19].

Normal healthy ageing leads to a variety of sensory and cognitive changes, including loss of sensory acuity [20–22], reduced processing speed [23], and impaired attentional and working memory processes [24, 25]. In multisensory perception, ageing has been associated with altered susceptibility to the sound-induced flash and McGurk illusions [26–30]; these age differences may be caused by various computational or neural mechanisms, including changes in sensory acuity, prior binding tendency, and attentional resources (for further discussion see [31]). By contrast, older adults perform in a way that is comparable to their younger counterparts on audiovisual integration of spatial signals (as indexed by the spatial ventriloquist illusion) [32, 33]. They arbitrate between sensory integration and segregation effectively, and weight signals in a way that is consistent with normative Bayesian Causal Inference. However, they sacrifice response speed to maintain this audiovisual localisation accuracy [32].

This raises the question of *how* older adults preserve audiovisual integration and spatial localisation performance, albeit with slower response times, in these intersensory selective attention paradigms. One possibility is that older adults rely on the same neural systems as younger adults, but neural processing takes longer to obtain comparable levels of accuracy. For instance, older adults may accumulate noisier audiovisual evidence for longer until they reach a decision threshold and commit to a response, as recently suggested by computational modelling of behavioural data [32]. Further, older adults may exert more top-down attentional control during this accumulation process to attenuate internal sensory noise. This longer, and more attentionally demanding, evidence accumulation would be reflected in increased BOLD responses, particularly in higher-order association cortices (e.g. parietal cortices) for older relative to younger adults. Critically, however, because the regional BOLD response reflects the accumulated neural activity, the information about task-relevant variables that can be decoded from it should be comparable in both age groups.

Alternatively, older adults may engage additional cortical regions to compensate for encoding deficits in the brain regions that are activated by both age groups. In this case, we would expect age differences not only in the magnitude of the regional BOLD responses, but also in their information content. In this latter case, the additional brain activations would encode more task-relevant information in older than in younger participants.

To adjudicate between these two hypotheses, we presented healthy younger and older participants with synchronous audiovisual signals at varying degrees of spatial disparity in a spatial ventriloquist paradigm. In an auditory selective attention task, participants reported the location of the auditory signal, whilst ignoring the task-irrelevant visual signals (which were spatially congruent or incongruent). Using multivariate pattern analysis (MVPA), we first tested whether the age groups similarly combined audiovisual signals into spatial representations along the dorsal visual and auditory spatial processing hierarchies that have previously been shown to be engaged in this task [10, 13]. Whole-brain univariate analyses then delineated neural systems that were commonly activated by both age groups during the task, as well as systems showing greater activations in older participants. Finally, using multivariate Bayesian decoding (MVB) [34], we assessed whether the regions with greater activation in older adults encoded critical stimulus information (such as visual and auditory location or their spatial relationship) to a greater degree in older than younger adults.

## Results

### Auditory spatial classification performance for older and younger adults

We assessed whether ageing impacts the precision with which older adults encode sound location. In a spatial left-right classification task (outside the scanner), older and younger adults were presented with unisensory auditory stimuli sampled randomly from 10 possible spatial locations along the azimuth. We fitted psychometric functions to the proportions of perceived ‘right’ responses individually for each participant and compared the JND (just-noticeable difference; i.e. spatial reliability or sensitivity) and PSE (point of subjective equality; left/right bias) between older and younger participants in two-sample *t* tests. We observed no significant differences in spatial precision or left/right bias between age groups; only a non-significant trend of larger JNDs (lower auditory spatial reliability) was evident in older adults: JND *t*(30) = 1.532, *p* = .136, *d* = 0.542; PSE *t*(30) = 0.527, *p* = .602, *d* = 0.186. This suggests comparable localisation performance for older and younger participant groups in an unspeeded auditory spatial classification task.

### Audiovisual integration behaviour for older and younger adults (inside the scanner)

In the main experiment inside the scanner, participants were presented with synchronous auditory and visual signals at the same (i.e. congruent) or opposite (i.e. incongruent) locations sampled from four possible spatial locations (-15°, -5°, 5°, or 15° visual angle) along the azimuth. The experimental design thus conformed to a 4 (auditory location: -15°, -5°, 5°, or 15° azimuth) x 3 (sensory context: unisensory auditory, audiovisual congruent, audiovisual incongruent) factorial design (see Fig 1B). On each trial, participants reported their perceived sound location as accurately as possible by pressing one of four spatially corresponding buttons with their right hand. For behavioural analysis, we pooled over hemifield and entered observers’ response accuracy and reaction times into a 2 (eccentricity: small ± 5° vs. large ± 15°) x 3 (sensory context: unisensory auditory, audiovisual congruent, audiovisual incongruent) x 2 (age group: younger, older) mixed ANOVA. For localisation accuracy, this mixed ANOVA identified significant main effects of eccentricity and sensory context (see Table 1). Further, a small three-way (eccentricity x sensory context x age) interaction was observed, reflecting a slightly stronger visual influence on perceived sound location in older adults for audiovisual stimuli at large spatial disparities (see right panel of Fig 1C). This stronger audiovisual crossmodal bias in older adults was not observed in previous research that was performed outside the scanner [32]. This small discrepancy between studies may be explained by the adverse listening conditions inside the scanner that make it more difficult for observers to reliably arbitrate between sensory integration and segregation, even at large spatial disparities (see [9]). No other significant effects were observed.

**Fig 1.**
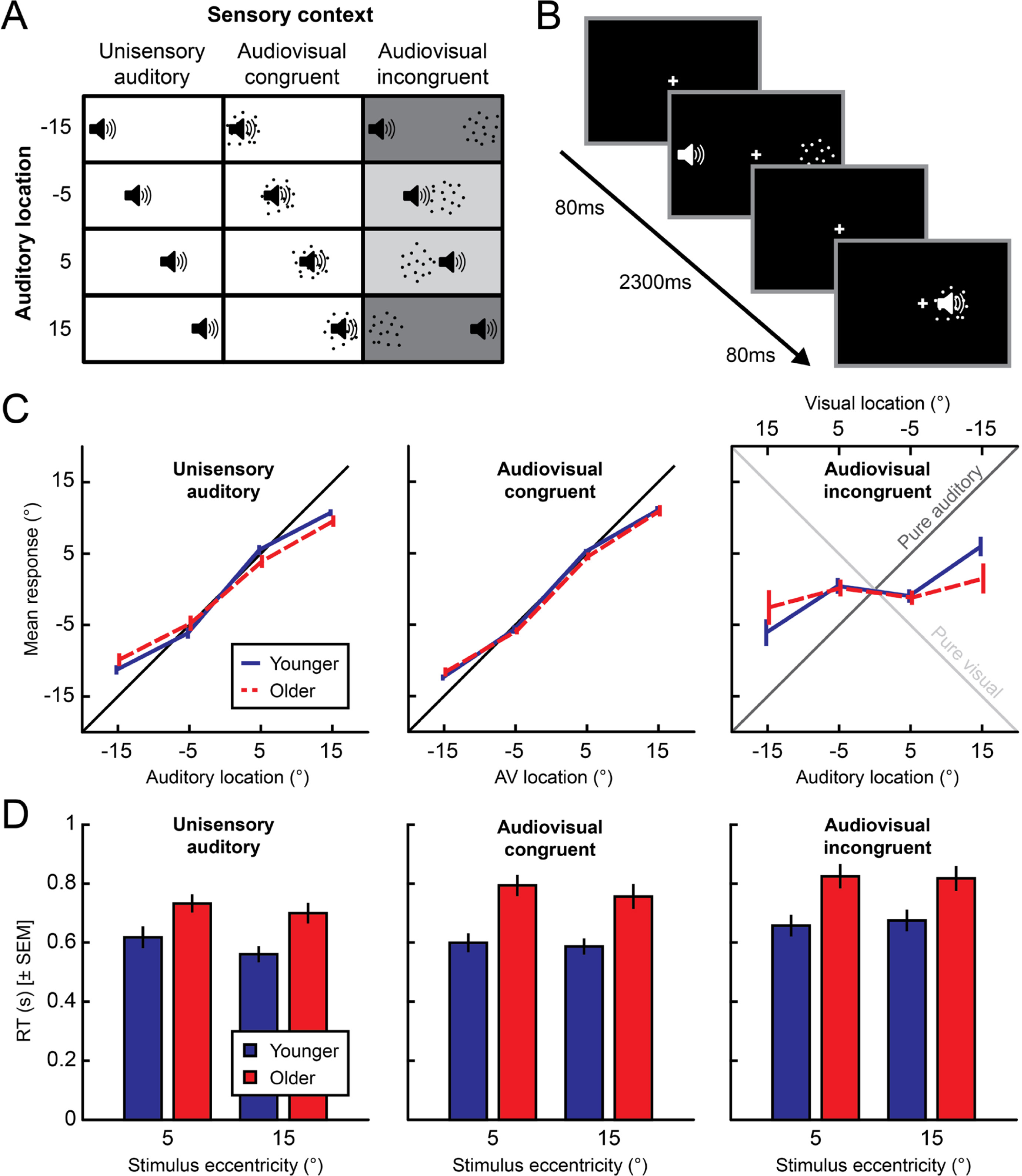
Experimental design and behavioural results. (A and B) The experiment conformed to a 4 (auditory location) × 3 (sensory context: unisensory auditory, audiovisual congruent, audiovisual incongruent) factorial design. Auditory (white noise bursts) and visual signals (cloud of dots) were sampled from four possible azimuthal locations (-15°, -5°, 5°, or 15°). Auditory and visual stimuli were presented either at same (congruent) or opposite spatial locations (incongruent). Participants reported their perceived location of the sound. (C) Across-participants mean (± *SEM*) perceived sound locations as a function of the true sound location (*x* axis). Older and younger adults showed comparable central biases (i.e. deviations from the identity line) for unisensory and audiovisual congruent stimuli. For spatially incongruent stimuli, older adults showed a slightly stronger spatial bias in their perceived sound location towards the location of the incongruent visual signal. (D) Behavioural response times (pooled over left and right hemifields; across-participants means of condition-specific medians). Participants responded more slowly to audiovisual incongruent relative to audiovisual congruent and auditory-only stimuli. Older adults were significantly slower in all conditions, but this did not interact with any other factor.

**Table 1.** Results of mixed ANOVAs on mean auditory localisation responses and median reaction times during the spatial ventriloquist task (inside the scanner).

| | *df* | | *F* | *p* | $\eta_p^2$ |
| --- | --- | --- | --- | --- | --- |
|  | effect | error |  |  |  |
| *Mean localisation responses* |  |  |  |  |  |
| Eccentricity | 1 | 30 | 262.844 | < **.001** | .898 |
| Eccentricity x Age | 1 | 30 | 3.970 | .055 | .117 |
| Sensory context | 1.144 | 34.335 | 51.117 | < **.001** | .630 |
| Sensory context x Age | 1.144 | 34.335 | 0.646 | .447 | .021 |
| Eccentricity x Sensory context | 1.204 | 36.111 | 2.045 | .159 | .064 |
| Eccentricity x Sensory context<br>x Age | 1.204 | 36.111 | 5.344 | **.021** | .151 |
| Age | 1 | 30 | 2.189 | .149 | .068 |
| *Median reaction times* |  |  |  |  |  |
| Eccentricity | 1 | 30 | 3.261 | .081 | .098 |
| Eccentricity x Age | 1 | 30 | 0.119 | .733 | .004 |
| Sensory context | 2 | 60 | 34.145 | < **.001** | .532 |
| Sensory context x Age | 2 | 60 | 3.010 | .057 | .091 |
| Eccentricity x Sensory context | 1.575 | 47.236 | 6.095 | **.008** | .169 |
| Eccentricity x Sensory context<br>x Age | 1.575 | 47.236 | 1.944 | .162 | .061 |
| Age | 1 | 30 | 10.914 | **.002** | .267 |

Degrees of freedom Greenhouse-Geisser corrected for non-sphericity where applicable.

The corresponding 2 x 3 x 2 mixed ANOVA of participants’ median reaction times (inside the scanner) revealed main effects of age and sensory context as well as an interaction between sensory context and eccentricity (see Table 1). Older adults were overall slower than younger adults. Participants responded fastest to unisensory auditory stimuli, slower to audiovisual congruent stimuli, and slowest to audiovisual incongruent stimuli. The longer response times for audiovisual congruent compared to unisensory auditory stimuli is a surprising finding that may again be explained by the causal uncertainty invoked by the competing scanner noise. Because unisensory auditory, congruent audiovisual, and incongruent audiovisual stimuli were presented intermixed, observers needed to infer whether audiovisual signals came from the same source and should thus be integrated. Causal inference becomes more challenging in adverse listening situations, placing extra attentional demands on our audiovisual trials that may outweigh any multisensory benefit. Further, as indicated by the significant interaction between eccentricity and sensory context, observers were slower to respond to more centrally than peripherally presented sounds, particularly in the unisensory auditory context. None of these effects significantly interacted with age, however. See Table 1 for detailed results of sound localisation responses and response times. In summary, while older adults were substantially slower than younger adults across all conditions, their auditory localisation performance was largely comparable to their younger counterparts.

### fMRI results

We used fMRI to assess the commonalities and differences in the neural systems underlying audiovisual spatial processing between younger and older adults, in three steps. First, we characterised how younger and older participants integrate auditory and visual information into spatial representations along the dorsal audiovisual processing hierarchies, using support vector regression. Second, we identified commonalities and differences in regional BOLD responses for older and younger adults, using mass-univariate fMRI analyses. Third, we investigated whether age-related BOLD-response increases encode critical stimulus information (such as visual and auditory location or their spatial relationship) to a greater degree in older than younger adults, using multivariate Bayesian decoding (MVB) [34].

### Decoding audiovisual spatial estimates using support vector regression

Fig 2 shows the spatial locations decoded with support vector regression from regional BOLD-response patterns for unisensory auditory, congruent audiovisual, and incongruent audiovisual incongruent stimuli along the dorsal auditory and visual processing hierarchies (see also Tables 2 and 3). As previously reported for younger populations [10, 13], primary auditory area A1 and “higher-level” auditory area planum temporale (PT) encoded mainly the sound location, while “low-level” visual areas V1-V3, posterior intraparietal sulcus (IPS 0-2) and, anterior intraparietal sulcus (IPS 3-4) represented the visual location.

**Fig 2.**
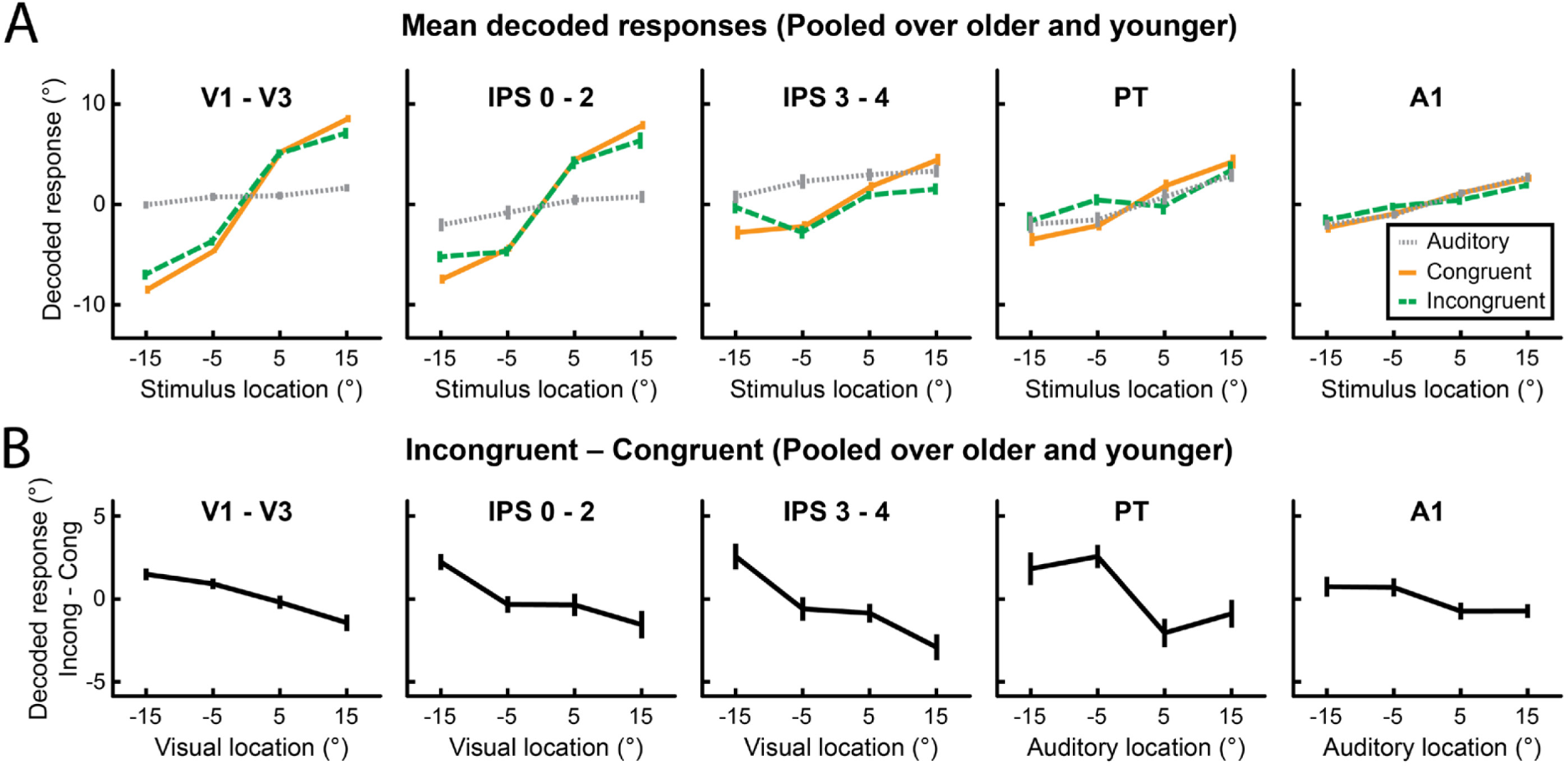
fMRI multivariate decoding results (support vector regression) pooled over age groups. (A) Across-participants mean (±1 *SEM)* decoded spatial locations for unisensory auditory (grey), audiovisual congruent (orange), and audiovisual incongruent (green) stimuli. (B) Difference between decoded stimulus locations for audiovisual incongruent relative to audiovisual congruent stimuli. Results for five ROIs are shown: visual regions (V1 - V3); posterior intraparietal sulcus (IPS 0 - 2); anterior intraparietal sulcus (IPS 3 - 4); planum temporale (PT); and primary auditory cortex (A1). Note that the *x* axis is labelled according to each region’s dominant sensory modality (i.e. visual location for V1-3 and IPS, auditory location for PT and A1) to allow for easier comparison between conditions and regions.

**Table 2.**
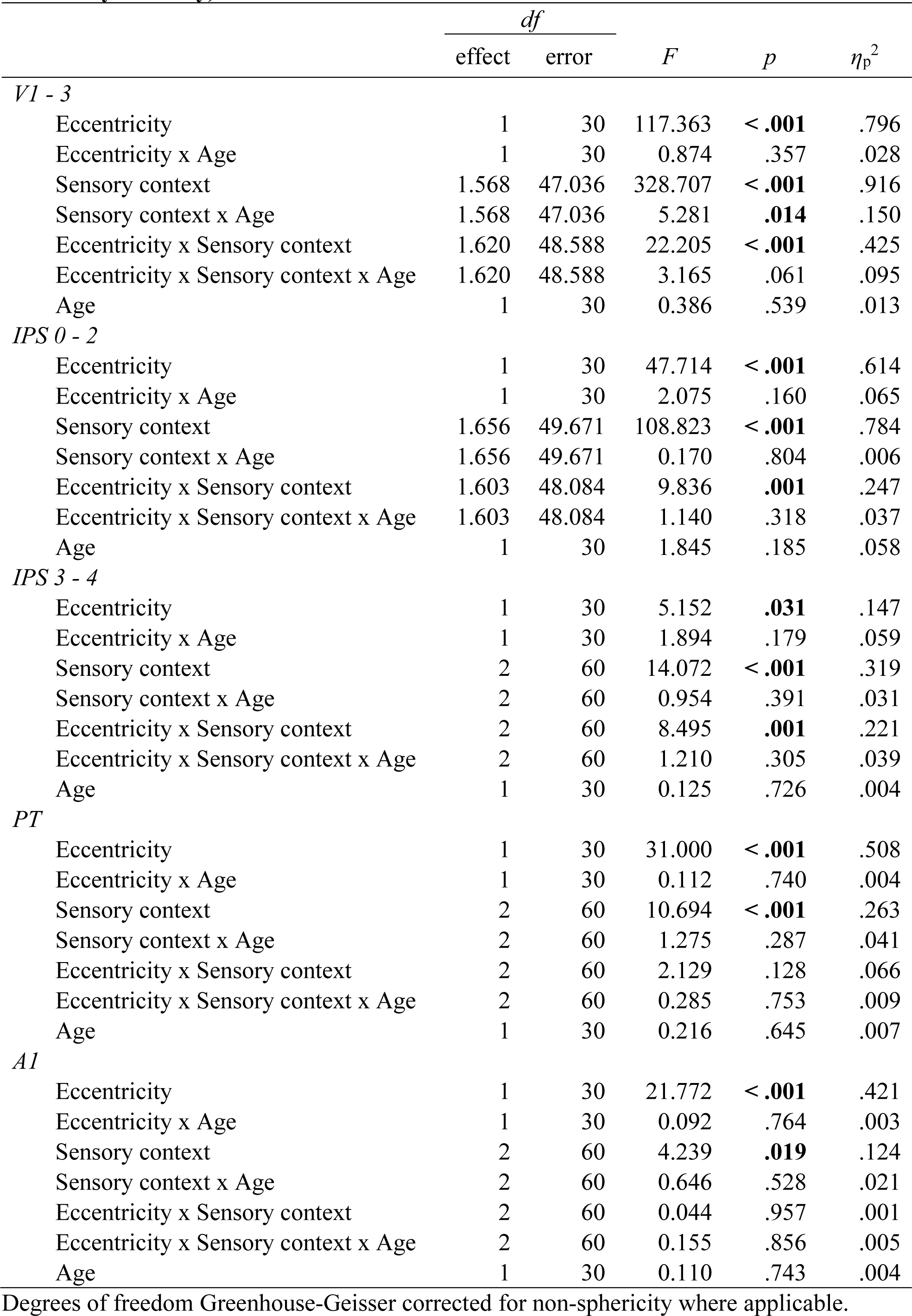
Results of ANOVAs on SVR decoded responses in five ROIs (including unisensory auditory).

Importantly, the decoding profiles differed for congruent and incongruent audiovisual stimuli in all regions. In auditory area PT, incongruent visual inputs biased auditory spatial encoding mainly at small spatial disparities (i.e. a “neural ventriloquist effect”). These crossmodal biases broke down at large spatial disparities, when the brain infers that two signals come from different sources - thereby mirroring the integration profile observed at the behavioural level. Surprisingly, in visual areas we observed an influence of a displaced sound on the decoded spatial location mainly at large spatial disparities. This pattern may be explained by the fact that at small spatial disparities, observers experience a ventriloquist illusion and thus perceive the sound shifted towards the visual signal. By contrast, at large spatial disparities (when observers are less likely to experience a ventriloquist illusion), a displaced sound from the opposite hemifield biases the spatial encoding in visual cortices via mechanisms of top-down attention. As previously reported [10, 13], these crossmodal interactions increased across the cortical hierarchy, being more pronounced in IPS and PT than in early visual and auditory cortices.

These impressions were confirmed statistically by the 2 (eccentricity: small, large) x 3 (sensory context: unisensory auditory, audiovisual congruent, audiovisual incongruent) x 2 (age: younger, older) mixed ANOVAs of the decoded spatial estimates, separately for each region of interest (ROI) along the visual and auditory processing hierarchy (Table 2). We observed main effects of stimulus eccentricity for all ROIs, confirming that all regions encoded information about the location of the stimuli. Intriguingly, main effects of sensory context were also present in all ROIs, suggesting that even putatively unisensory regions held at least some information about whether a visual stimulus was present and/or its spatial congruence with the sound. We confirmed that these sensory context effects were not driven entirely by differences between unisensory auditory vs. audiovisual stimuli: a follow-up ANOVA that excluded the unisensory condition, so 2 (eccentricity: small, large) x 2 (congruency: congruent vs incongruent) x 2 (age group: older vs. younger), revealed a significant main effect of congruency for all ROIs, and a significant congruence x eccentricity interaction in areas V1 – V3, IPS 0 – 2, and IPS 3 – 4 (for detailed results see tables in Supporting Information).

Crucially, however, age had almost no effect on the locations decoded from the activation patterns along the auditory and visual spatial processing hierarchies (see Fig 3 and Tables 2 and 3). We observed a single significant age-related effect across all ANOVAs: an age x sensory context interaction selectively in visual areas V1 to V3. However, in the follow-up ANOVA that excluded the unisensory auditory condition, the age x congruency interaction was not significant, and independent-samples *t* tests comparing the age groups in all conditions only revealed an age difference for unisensory stimuli at large eccentricities (*t*(30) = 2.623, *p* = .014 (uncorr.), *d* = 0.927; see leftmost panel of Fig 3A). Collectively, these results compellingly demonstrate that younger and older adults similarly combine auditory and visual signals into spatial representations in regions along the auditory and visual processing hierarchies.

**Fig 3.**
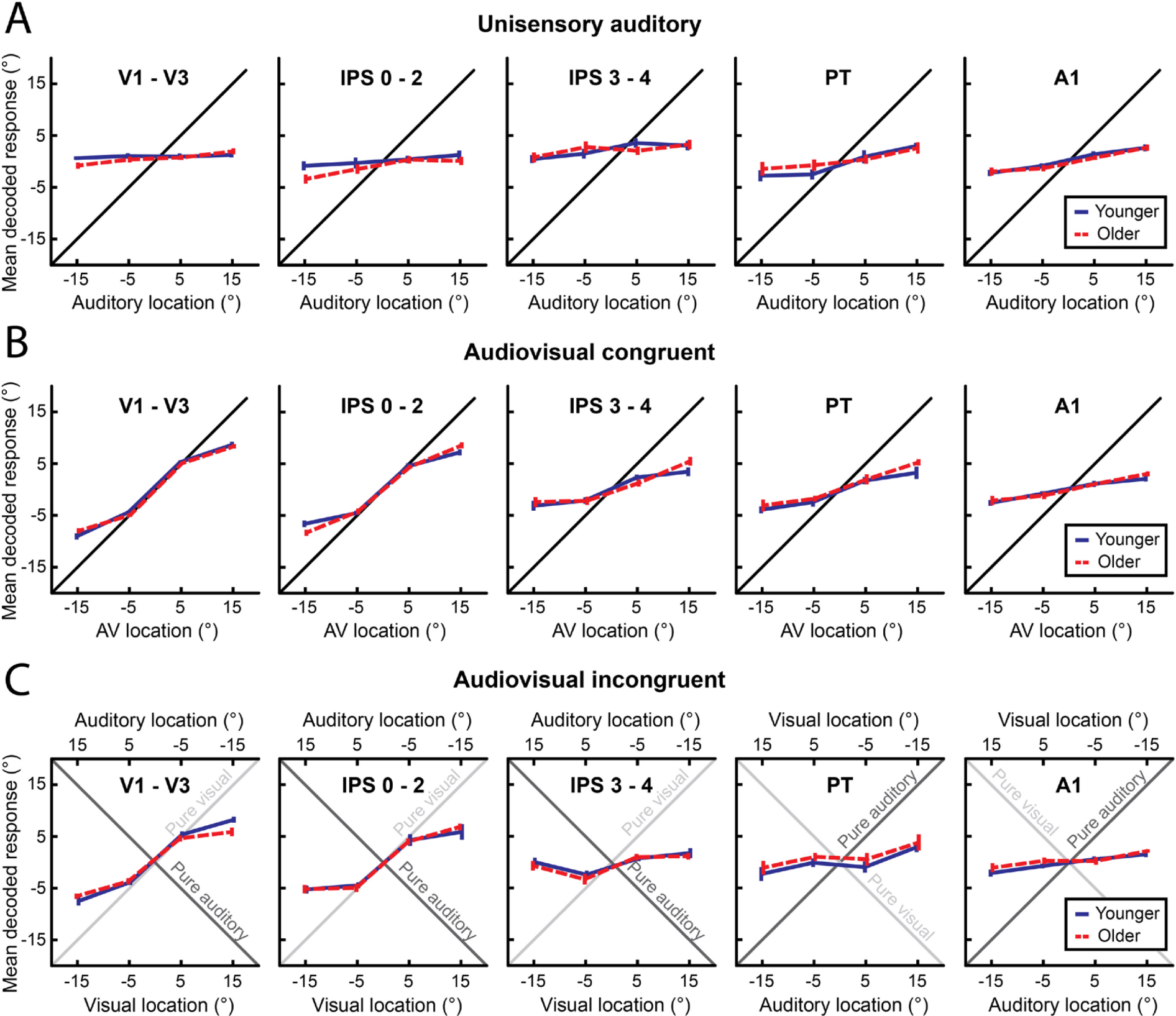
fMRI multivariate decoding results (support vector regression) separately for older and younger adults. Across-participants mean (±1 *SEM)* decoded spatial locations for younger (blue) and older (red) participants for (A) unisensory auditory, (B) congruent audiovisual, and (C) incongruent audiovisual stimuli. Results for five ROIs are shown: visual regions (V1 - V3); posterior intraparietal sulcus (IPS 0 - 2); anterior intraparietal sulcus (IPS 3 - 4); planum temporale (PT); and primary auditory cortex (A1). Note that for incongruent conditions the location of stimuli in the region’s dominant sensory modality is plotted on the lower *x* axis (i.e. visual location for V1-3 and IPS, auditory location for PT and A1) to allow for easier comparison between conditions and regions.

### Conventional mass-univariate GLM analysis

The above SVR analysis showed that regions along the auditory and visual spatial processing hierarchies integrate sensory signals into spatial representations similarly in both age groups. Using mass-univariate general linear model (GLM) analysis, we next investigated whether older and younger adults engage overlapping or partly distinct neural systems for audiovisual processing (i.e. all stimulus conditions > fixation). Moreover, we assessed the neural underpinnings of cognitive control and attentional operations that are critical for localising a sound when presented together with a spatially displaced visual signal (i.e. incongruent > congruent audiovisual stimuli; see Table 3, and Figs 4 and 5, for details).

**Fig 4.**
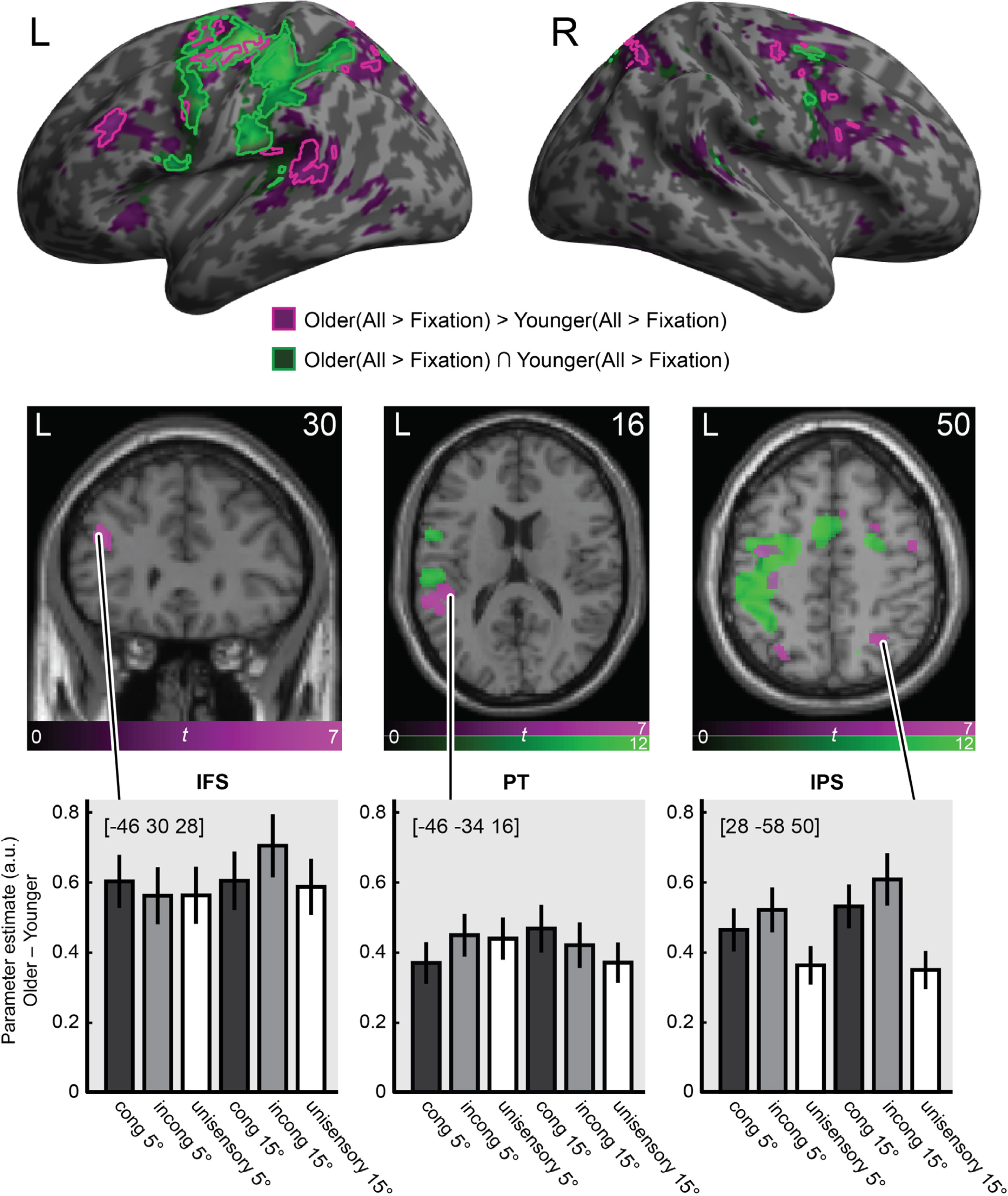
fMRI activation results for older and younger adults. Activations for all stimuli (i.e. pooled over auditory, audiovisual congruent and incongruent) relative to fixation are rendered on an inflated canonical brain (top row) and coronal/transverse sections (middle row). Green = conjunction over both age groups (All_Older_ > Fixation_Older_) ∩ (All_Younger_ > Fixation_Younger_). Purple = age related activation increases (All_Older_ > Fixation_Older_) > (All_Younger_ > Fixation_Younger_). For inflated brain: bright outlines = height threshold *p* < .05 whole-brain FWE-corrected. For visualisation purposes we also show activations at *p* < .001, uncorrected, as darker filled areas. Extent threshold *k* > 0 voxels). For brain sections, height threshold *p* < .05 whole-brain FWE-corrected. Bottom row: Bar plots show mean (± 1 *SEM*) age differences in parameter estimates (arbitrary units) for audiovisual congruent, audiovisual incongruent, and unisensory auditory stimuli at 5° and 15° eccentricities, pooled over left and right stimulus locations, at the indicated peak MNI coordinates. Three illustrative anatomical regions are shown: left inferior frontal sulcus [IFS], left planum temporale [PT], and right intraparietal sulcus [IPS].

**Fig 5.**
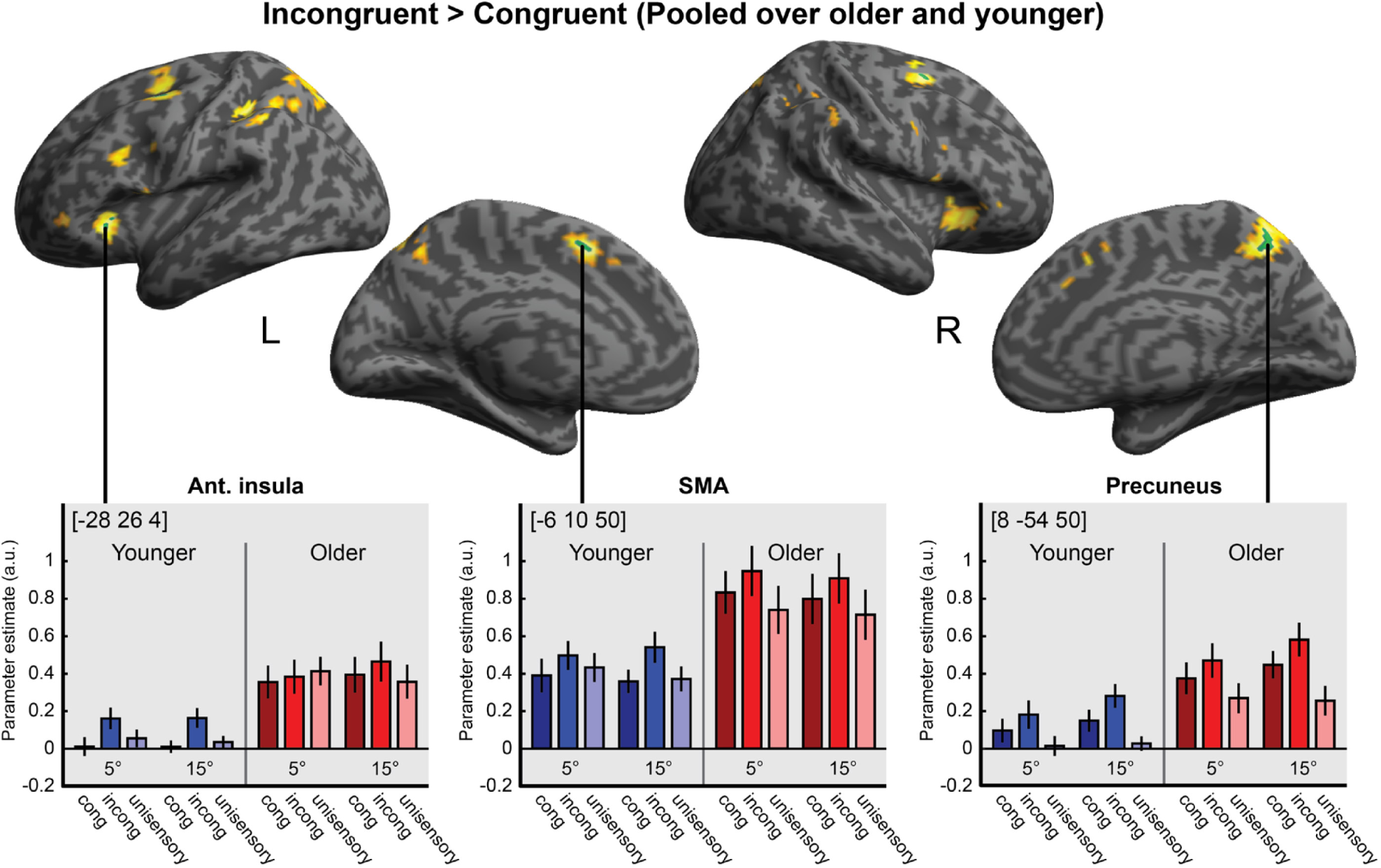
Activation increases for incongruent > congruent audiovisual stimuli. Activation increases for incongruent relative to congruent stimuli (pooled over age groups) are rendered on an inflated canonical brain. Green areas = height threshold *p* < .05, whole-brain FWE-corrected. For visualisation purposes we also show activations at *p* < .001, uncorrected, in yellow. Bar plots show parameter estimates (across-participants mean ± 1 *SEM*; arbitrary units) for congruent, incongruent, and unisensory stimuli at 5° and 15° eccentricities, pooled over left and right auditory (and in audiovisual conditions, visual locations), at the indicated MNI peak coordinates in three anatomical regions: left anterior insula, left supplementary motor area (SMA), and right precuneus.

**Table 3.** Mass univariate fMRI analysis – results.

| Region | Coordinates |  |  | z-score | p-value<br>(FWE*) |
| --- | --- | --- | --- | --- | --- |
| *O(All > Fixation) ∩ Y(All > Fixation)* |  |  |  |  |  |
| R. cerebellum | 22 | -54 | -24 | > 8 | < .001 |
| R. cerebellum | 6 | -62 | -16 | 6.9 | < .001 |
| R. cerebellum | 8 | -72 | -16 | 5.9 | < .001 |
| L. precentral gyrus | -36 | -20 | 64 | > 8 | < .001 |
| L. precentral sulcus | -32 | -4 | 58 | > 8 | < .001 |
| L. intraparietal sulcus | -46 | -34 | 42 | > 8 | < .001 |
| L. supplementary motor area | -4 | 0 | 56 | > 8 | < .001 |
| R. superior frontal sulcus | 24 | -2 | 50 | 5.7 | < .001 |
| L. thalamus | -14 | -18 | 6 | 5.4 | 0.002 |
| L. intraparietal sulcus | -18 | -68 | 54 | 5.4 | 0.002 |
| R. precentral gyrus | 52 | 4 | 42 | 5.1 | 0.005 |
| L. planum temporale | -40 | -36 | 10 | 5.1 | 0.007 |
| L. anterior insula | -30 | 18 | 8 | 5.0 | 0.009 |
| L. superior frontal gyrus | -16 | -6 | 68 | 5.0 | 0.011 |
| R. intraparietal sulcus | 14 | -66 | 52 | 4.9 | 0.014 |
| R. superior temporal gyrus | 58 | -34 | 14 | 4.8 | 0.027 |
| *Incong > Cong (Pooled over age groups)* |  |  |  |  |  |
| R. precuneus | 8 | -54 | 50 | 5.2 | < .001 |
| L. supplementary motor area | -6 | 10 | 50 | 5.0 | < .001 |
| L. superior frontal sulcus | -26 | 6 | 58 | 5.0 | < .001 |
| L. superior frontal sulcus | -26 | -2 | 48 | 4.9 | < .001 |
| L. anterior insula | -28 | 26 | 4 | 5.0 | < .001 |
| R. superior frontal sulcus | 24 | 2 | 54 | 4.8 | < .001 |
| R. anterior insula | 32 | 26 | -4 | 4.8 | < .001 |
| L. superior frontal sulcus | -30 | -2 | 62 | 4.7 | < .001 |
| *O(All > Fixation) > Y(All > Fixation)* |  |  |  |  |  |
| L. inferior frontal sulcus | -46 | 30 | 28 | 7.3 | < .001 |
| L. precentral gyrus | -38 | -8 | 54 | 6.6 | < .001 |
| L. supplementary motor area | -8 | -8 | 64 | 6.3 | < .001 |
| L. superior frontal sulcus | -20 | -8 | 56 | 5.8 | < .001 |
| L. superior temporal gyrus | -60 | -40 | 12 | 5.8 | < .001 |
| L. planum temporale | -46 | -34 | 16 | 5.6 | .001 |
| L. supramarginal gyrus | -50 | -44 | 22 | 5.4 | .001 |
| R. intraparietal sulcus | 28 | -58 | 50 | 5.6 | .001 |
| R. precuneus | 12 | -62 | 62 | 5.5 | .001 |
| R. intraparietal sulcus | 24 | -62 | 56 | 5.0 | .011 |
| R. precentral sulcus | 48 | -4 | 52 | 5.6 | .001 |
| R. supplementary motor area | 8 | 18 | 46 | 5.5 | .001 |
| R. inferior frontal sulcus | 36 | 2 | 36 | 5.4 | .002 |
| L. precuneus | -10 | -64 | 58 | 5.3 | .002 |
| L. intraparietal sulcus | -26 | -70 | 50 | 5.2 | .004 |
| R. superior frontal sulcus | 26 | -6 | 56 | 5.2 | .004 |
| R. supplementary motor area | 10 | 6 | 56 | 5.2 | .005 |
| R. superior frontal sulcus | 26 | 6 | 54 | 5.2 | .005 |
| L. precentral sulcus | -46 | 6 | 34 | 5.1 | .007 |
| L. precentral sulcus | -50 | -8 | 46 | 5.0 | .012 |
| L. intraparietal sulcus | -28 | -54 | 46 | 4.9 | .014 |
| L. superior temporal pole | -52 | 14 | -4 | 4.9 | .018 |
| R. inferior frontal sulcus | 38 | 14 | 26 | 4.9 | .019 |
| L. intraparietal sulcus | -24 | -62 | 58 | 4.8 | .031 |
| L. intraparietal sulcus | -44 | -40 | 34 | 4.7 | .037 |
| L. anterior insula | -30 | 24 | 0 | 4.7 | .047 |

\**p* values whole-brain corrected for family-wise errors at the voxel level.

### Effects of stimuli and task relative to fixation

A conjunction analysis over age groups revealed stimulus-induced activations in a widespread neural system encompassing key areas of the auditory spatial processing hierarchy such as left planum temporale, extending into left inferior parietal lobe and intraparietal sulci bilaterally *(All_Older_ > Fixation_Older_) ∩ (All_Younger_ > Fixation_Younger_)* [35, 36]. At a lower threshold of significance, we also observed stimulus-induced activations in the right hemisphere from right planum temporale into inferior parietal lobe and bilateral insulae. Moreover, we observed common activations related to response selection and motor processing in left precentral gyrus/sulcus and right cerebellum.

Next, we identified regions with greater activations for older relative to younger adults by testing for the interaction *(All_Older_ > Fixation_Older_) > (All_Younger_ > Fixation_Younger_).* We observed activation increases for older adults in dorsolateral prefrontal cortices along the inferior frontal sulcus. Interestingly, increased activations for older adults were often found adjacent to the regions that were commonly activated for both groups. For instance, we observed greater activations in the lateral plana temporalia extending into more posterior superior temporal cortices. Likewise, the parietal activations extended from the activation clusters observed for both age groups more posteriorly. Moreover, older adults showed increased activations in the inferior frontal sulcus, a region previously implicated in cognitive control of audiovisual processing tasks [37, 38]. In summary, older adults showed increased activations relative to younger adults along the spatial auditory pathways from temporal to parietal and frontal cortices.

The opposite contrast *(All_Younger_ > Fixation_Younger_) > (All_Older_ > Fixation_Older_)* revealed no activations that were significantly greater in the younger age group.

### Effects of audiovisual spatial incongruency

Consistent with previous research [14,37–39], incongruent relative to congruent audiovisual stimuli increased activations in a widespread attentional and cognitive control system including medial and lateral posterior parietal cortices, inferior frontal sulcus and bilateral anterior insulae (i.e. *Incong > Cong,* pooled over age groups). However, none of these incongruency effects interacted with age group after whole-brain correction *(Incong_Older_ > Cong_Older_) > (Incong_Younger_ > Cong_Younger_)* or *(Incong_Younger_ > Cong_Younger_) > (Incong_Older_ > Cong_Older_)*.

### Multivariate Bayesian decoding

The activation increases for older relative to younger adults raise the critical question of whether/how they contribute to sound localisation performance in older adults. Do these activations encode information about task-relevant variables such as stimulus location or audiovisual congruency, thereby enabling older adults to maintain auditory localisation accuracy? To address this question, we used model-based multivariate Bayesian decoding, which allows one to compare the ability of activation patterns in different brain regions to predict target variables. Specifically, we compared the predictive ability of three candidate ROIs: i. the regions activated jointly by older and younger adults [O∩Y], ii. the regions activated more by older than younger adults [O>Y], and iii. the union of the two [O>Y ∪ O∩Y]. We computed multivariate Bayesian decoding models separately for four target variables that relate to stimulus properties such as visual location, auditory location and spatial disparities (VisL ≠ VisR, AudL ≠ AudR, Incong5 ≠ Cong5, and Incong15 ≠ Cong15). To match the number of features across ROIs we limited each model to the most significant 1000 voxels in each ROI (see Materials and Methods for details). Summed over participants, log model evidence was greater for the [O>Y] than for the [O∩Y] ROI for all target variables, suggesting that older participants show greater activations in regions that encode stimulus-relevant information in both age groups. Indeed, as shown in Fig 4, the age-related activation increases are found particularly in planum temporale and parietal cortices that have previously been shown to be critical for encoding spatial information about auditory and visual stimuli and their spatial congruency [10,40,41]. Moreover, the union model [O>Y ∪ O∩Y] outperformed the more parsimonious models [O∩Y] and [O>Y] for each of the target variables. Bayesian model selection indicated that the protected exceedance probability was above 0.81 for the union model across all target variables in both age groups (see Fig 6).

**Fig 6.**
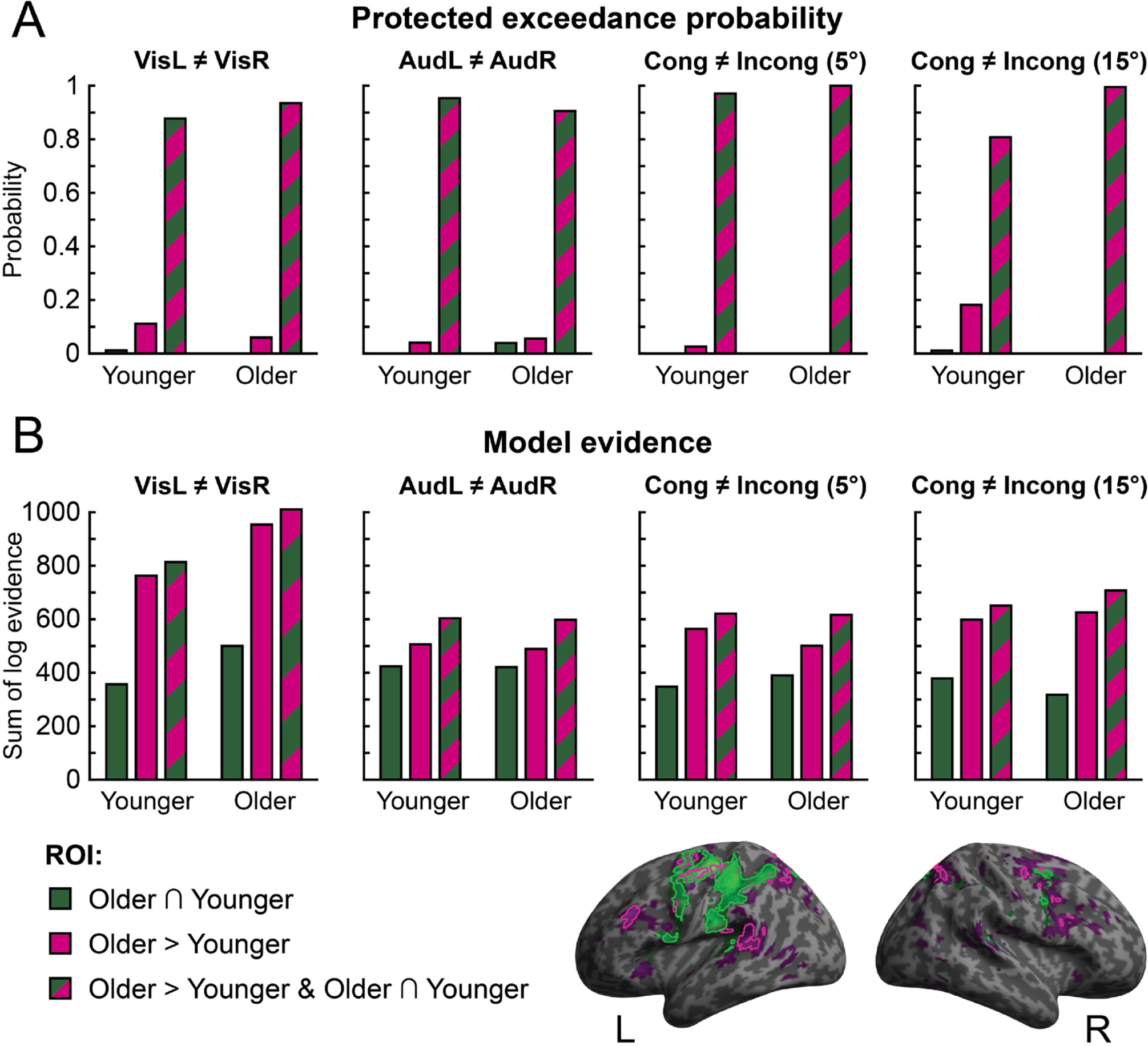
Results of multivariate Bayesian decoding analysis (MVB). Comparison of three ROIs ([O∩Y], [O>Y] or union of both: [O>Y ∪ O∩Y]) in their ability to predict stimulus related target variables: visual location, auditory location, congruent/incongruent at 5° and congruent/incongruent at 15°. (A) Log model evidences, summed over participants, are shown separately for each target variable and age group. (B) Random-effects Bayesian model comparison across the three ROIs, separately for each target variable and age group: protected exceedance probabilities for each ROI and target variable are shown.

These model comparison results collectively show that, in both age groups, the regions with greater activations in older adults [O>Y] encode significant information about task-relevant variables that is complementary to the information encoded in regions commonly activated by younger and older adults [O∩Y].

Next, we asked whether this increase in stimulus and task-relevant information for [O>Y] regions is more prevalent or important in older adults, as they show more activations in these regions. To address this question, we assessed whether the union [O>Y ∪ O∩Y] relative to the more parsimonious models [O∩Y] and [O>Y] won more frequently in the older age group. Contrary to this conjecture, there were no significant age differences in the frequency with which the union model was the winning model for predicting any of the four target variables (*χ*² tests of association, *p* > .05).

To further explore possible age differences, we investigated the relative contributions of the three ROIs to the encoding of task-relevant variables in older and younger participants by entering the difference in log model evidence for the union [O>Y ∪ O∩Y] ROI relative to the O∩Y ROI for each older and younger participant into Mann-Whitney U tests, separately for each of the four target variables. None of these tests revealed any significant differences between age groups across the VisL ≠ VisR, AudL ≠ AudR, and Incong5 ≠ Cong5 target variables, *p* > .05. Only for the Incong15 ≠ Cong15 target variable did we observe a small, non-significant trend for a greater “boost” in model evidence for the union [O>Y ∪ O∩Y] ROI, relative to the O∩Y ROI, for older adults compared to younger adults, *U* = 69.000, *p* = .052, one tailed (Bonferroni corrected for the comparisons across the four target variables).

Taken together, these results suggest that task-relevant information is encoded in each of the ROIs and, in particular, in the ROIs that are more strongly activated by older adults [O>Y], suggesting that older adults boost activations in brain regions that are critical for task-performance and encoding stimulus-relevant information. Further, the information encoded in the conjunction [O∩Y] and the ‘greater activation’ [O>Y] ROIs were not redundant but at least partly complementary, so that the union ROI [O>Y ∪ O∩Y] outperformed both of those more parsimonious models. Crucially, however, this was true for both older and younger adults. Likewise, the additional information gained by adding the ‘greater activation’ [O>Y] ROI to the conjunction [O∩Y] ROI was comparable in both age groups. These results suggest that older adults show increased activations in brain areas that are important for encoding stimulus- and task-relevant information to match the encoding capacities of their younger counterparts.

## Discussion

Healthy ageing leads to deficits in sensory processing and higher-order cognitive mechanisms. Nevertheless, older adults have been shown to maintain the ability to appropriately integrate and segregate audiovisual signals to aid stimulus localisation [32, 42]. The present study investigated the neural mechanisms that support this maintenance of performance.

Consistent with previous research [20,32,42,43], our behavioural results suggest that older adults were largely able to maintain spatial localisation accuracy for unisensory auditory and congruent audiovisual stimuli, but took substantially longer to respond than their younger counterparts. For spatially incongruent audiovisual stimuli we observed small but significant differences between the age groups. Specifically, at the larger (30°) spatial disparity, older adults’ sound localisation responses were more biased towards the location of the spatially conflicting visual stimulus. These stronger audiovisual spatial biases were not observed in previous behavioural research [32, 42], and we suggest that they result from the greater attentional resources that are needed to arbitrate between integration and segregation of audiovisual signals in the noisy environment of the MRI scanner. Background noise reduces a target sound’s signal-to-noise ratio, increasing the attentional resources required to identify and locate it, particularly in the presence of a highly salient and incongruent visual distractor (as in our large audiovisual disparity condition). As argued in a recent review [31], the greatest effects of ageing on multisensory integration are often found in situations of high attentional demand featuring, for example, noise or distractor signals (see e.g. [44–46]). Future behavioural research could further explore this hypothesis by assessing the effects of ageing on spatial localisation in a ventriloquist task under various degrees of background noise.

At the neural level, our multivariate analysis showed that audiovisual interactions increase progressively across the cortical hierarchy, as previously shown in human neuroimaging and neurophysiology studies [10,13,14,47–49]. Primary auditory cortices (A1) encoded primarily the location of the auditory component of the stimuli, and early visual cortices (V1 – V3) mainly that of the visual component, but small significant effects of sensory context and even audiovisual spatial congruency were observed even in these primary sensory areas. Again, these findings align nicely with a wealth of studies showing audiovisual interaction effects in primary sensory cortices [39,50–53]. Interestingly, a displaced visual stimulus biased the spatial encoding mainly at *small* spatial disparities in planum temporale, thereby mirroring the profile of crossmodal biases observed at the behavioural level. By contrast, a displaced auditory stimulus biased the spatial encoding mainly at *large* spatial disparities in visual cortices. The latter suggests that the crossmodal biases on spatial representations decoded from visual cortices arise mainly from top-down, possibly attentional, influences. At small spatial disparities the perceived location of the less spatially reliable sound is shifted towards the visual location, and thus does not affect spatial encoding in visual cortices. At large spatial disparities, audiovisual integration is attenuated or even abolished, so a spatially displaced sound may exert top-down attentional influences on the activation patterns in visual cortices.

Critically, however, none of these effects depended on age. Fig 3 shows that decoded stimulus locations were near identical in older and younger adults for unisensory auditory, congruent audiovisual, and incongruent audiovisual stimuli in all regions of interest. These results suggest that healthy ageing does not significantly alter how the brain integrates audiovisual inputs into spatial representations along the auditory or visual cortical pathways.

Despite these remarkably similar decoding profiles across the auditory and visual hierarchies between the two age groups, we observed significantly greater BOLD responses across an extensive network of frontal, temporal, and parietal regions for older adults in the spatial localisation task. This is in line with previous work showing age-related increases in BOLD response, especially in frontal and parietal regions, in a wide variety of situations [54–57], including those that involve processing of complex multisensory stimuli [58]. In the present study, older adults showed greater activations in areas including superior temporal cortices (including plana temporalia), as well as inferior frontal sulci and intraparietal sulci. Some of these areas were adjacent to, or even partly overlapped with, those activated by both age groups (i.e. task-relevant activations above baseline were present in both groups, but were greater in older adults).

This dissociation between age-related increases in regional BOLD responses, and comparable decoding profiles along the audiovisual pathways, raises the question of what these activation increases contribute to task performance. What is their functional role? In this study we aimed to distinguish between two possible mechanisms. First, older adults may compensate for their noisier sensory inputs via top-down attentional mechanisms and longer accumulation of noisy evidence into a decision variable in higher order association areas such as frontoparietal cortices [59, 60]. Indeed, recent computational modelling of audiovisual spatial localisation responses has suggested that older adults maintain spatial localisation accuracy by accumulating noisier sensory information for longer until a decision threshold is reached, and a response elicited [32]. Longer and more protracted evidence accumulation would be reflected in greater BOLD responses [38] for older than younger adults; yet, the regions with greater activations in older adults [O>Y] would contribute similarly to encoding relevant stimulus-relevant information (e.g. spatial location, congruency) across both age groups.

Second, older adults may recruit additional areas to compensate for processing and representational encoding deficits in other regions. This idea has previously been suggested for a variety of scenarios in which older adults also showed increased activations [54,61,62] (though see also [56, 63]). In this latter case, we would expect that the age-related activation increases encode information about task-relevant variables more strongly in older than in younger adults.

To adjudicate between these two potential neural mechanisms, we applied multivariate Bayesian analysis to compare the information about auditory location and audiovisual congruency that is encoded in areas with (1) joint activations in both age groups [O∩Y], (2) increased activations in older adults [O>Y], and (3) the union of those two sets of regions [O>Y ∪ O∩Y]. As expected, all three sets of regions encoded task-relevant information about sound location and audiovisual spatial disparity. Further, model comparison indicated that the ‘increased activations model’ outperformed the conjunction model. Yet, the union model still outperformed both more parsimonious models that included only one set of regions. Collectively, these results suggest that older adults enhance activations in brain areas that are critical for encoding stimulus-relevant information and that these regions provide stimulus-relevant information that is distinct (i.e. not redundant) from the information provided by the brain areas with joint activations. Crucially, this was true for both younger and older adults. Further, the boost in explanatory power when adding the [O>Y] ROI was also comparable in older and younger adults. Collectively, these results support our first proposed mechanism: that older adults engage similar neural systems for audiovisual integration, but need to accumulate noisier sensory evidence for longer, and exert greater top-down attentional control to enable reliable neural encoding of stimulus-relevant information, thus maintaining spatial localisation accuracy. Put differently, because older adults accumulate noisy evidence for longer, we observe age-related activation increases in the frontoparietal system. Further, this longer evidence accumulation allows older adults to obtain spatial representations of comparable spatial precision as their younger counterparts, which is reflected in the comparable spatial decoding accuracy at the neural level and spatial localisation accuracy at the behavioural level across age groups.

In conclusion, older adults have longer response times and greater frontoparietal activations than their younger counterparts. Yet, despite differences in BOLD-response magnitude, the spatial and stimulus-relevant information encoded in these regions is comparable across the two age groups. This dissociation—between comparable response accuracy and information encoded in brain activity patterns across the two age groups, but age-related increases in response times and regional activations—suggests that older participants accumulate noisier sensory evidence for longer, to maintain reliable neural encoding of stimulus-relevant information and thus preserve localisation accuracy.

## Materials and Methods

### Participants

Twenty younger and twenty-nine older adults were initially recruited from participant databases for a behavioural screening session. Two older adults were excluded from the study due to the presence of MRI contraindications, three failed to score above 24 on the Montreal Cognitive Assessment [64], and one reported taking antidepressant medication. A further seven older, and three younger, adults were excluded for insufficient gaze fixation in the behavioural task (see below for details). One younger participant could not be contacted following the behavioural session. Therefore, 16 younger (mean age = 24.19, *SD* = 4.56, 10 female) and 16 older (mean age = 70.75, *SD* = 4.71, 12 female) adults took part in all three experimental sessions. Those 32 included participants had normal or corrected-to-normal vision, reported no hearing impairment, and were able to distinguish left from right sounds with a just-noticeable difference (JND) of below 10°. The study was approved by the University of Birmingham Ethical Review Committee. All participants gave informed consent and were compensated for their time in cash or research credits.

### Stimuli

Visual stimuli consisted of an 80ms flash of 20 white dots (diameter of 0.4° visual angle), whose locations were sampled from a bivariate Gaussian distribution with a standard deviation of 2.5° in horizontal and vertical directions, presented on a black background.

Auditory spatialised stimuli (80 ms duration) were created by convolving a burst of white noise (with 5 ms onset and offset ramps) with spatially specific head-related transfer functions (HRTFs) based on the KEMAR dummy head of the MIT Media Lab [65]. Sounds were generated independently for every trial and presented with a 5ms on/off ramp.

### Design and procedure (for main experiment inside the MRI scanner)

In a spatial ventriloquist paradigm, participants were presented with synchronous auditory and visual signals at the same or different locations. The auditory signal originated from one of four possible spatial locations (-15°, -5°, 5°, or 15° visual angle) along the azimuth. For any given auditory location, a synchronous visual signal was presented at the same spatial location (audiovisual congruent trial), at the symmetrically opposite location (audiovisual incongruent trial), or was absent (unisensory auditory trial). On each trial, observers reported the sound location as accurately as possible by pressing one of four spatially corresponding buttons with their right hand. Thus, our design conformed to a 4 (auditory location: -15°, -5°, 5°, or 15° azimuth) x 3 (sensory context: unisensory auditory, audiovisual congruent, audiovisual incongruent) factorial design (see Fig 1B), though for behavioural statistical analyses on performance accuracy and response times we pooled over hemifields and rearranged the conditions into a 2 (eccentricity: ± 5° or ± 15°) x 3 (sensory context: unisensory auditory, audiovisual congruent, audiovisual incongruent) factorial design (see below). Participants fixated a central cross (white; 0.75° diameter) throughout the experiment. Trials were presented with a stimulus onset asynchrony (SOA) of 2.3 s. To increase design efficiency, the activation trials were presented in a pseudorandomised fashion interleaved with 6.9 s fixation periods approximately every 20 trials. The experiment included 10 trials (per condition, per run) x 12 conditions x 11 five-minute runs (split over two separate days).

### Experimental setup

Stimuli were presented using Version 3 of the Psychophysics Toolbox [66], running on MATLAB 2014b on an Apple MacBook. Auditory stimuli were presented at approximately 75 dB SPL through Optime 1 electrodynamic headphones (MR Confon). Visual stimuli were back-projected by a JVC DLA-SX21E projector onto an acrylic screen, viewed via a mirror attached to the MRI head coil. The total viewing distance from eye to screen was 68cm. Participants responded using infrared response pads (Nata Technologies) held in the right hand.

### Behavioural testing session (outside the scanner)

Participants took part in a total of three experimental sessions on three separate days (one behavioural, two MRI). In the first (behavioural) session they underwent training and screening.

First, in a left/right forced-choice spatial classification task, participants were presented on each trial with an auditory stimulus randomly at one of ten locations between - 15° and 15° azimuth (-15°, -10°, -5°, -3°, -1°, 1°, 3°, 5°, 10°, 15°) and indicated via a two-choice button press whether they perceived the sound as coming from the left or right.

Second, they were trained to learn the mapping between the auditory locations (-15°, - 5°, 5°, and 15°) and the four corresponding buttons used in the main ventriloquist paradigm. Via a four-choice key press, participants localised a sound that was presented randomly from one of the four locations on each trial. Feedback was provided after each response: correct responses were rewarded with a green square presented at the correct/responded location; incorrect responses resulted in a red square presented at the responded location, followed by a green square presented at the correct location. Participants completed up to five 20-trial blocks, stopping early if localisation accuracy (i.e. correct button responses) reached 90% in any block.

Third, participants completed two blocks of the spatial ventriloquist paradigm used during the two MRI scanning sessions. During these blocks, the scanner noise recorded from an fMRI sequence was played over speakers at a level that approximately matched that experienced in the scanner (after adjustment for headphone attenuation). Analyses on data from this session are included in the Supporting Information.

Because many older adults find remaining still for extended periods of time challenging and painful, we were unable to perform reliable eye tracking during fMRI scanning due to the associated extra setup and calibration times. Therefore, to minimise the possibility of eye movement confounds in the fMRI data, we instead screened participants beforehand for their ability to maintain central fixation during the task. Throughout the two blocks of the ventriloquist paradigm, participants’ eye movements were recorded via a Tobii EyeX eye tracker. A custom MATLAB script was used to remove blinks and identify saccades. For each participant, the peak location (i.e. furthest from fixation) of every recorded saccade was entered as the outcome variable in a linear regression analysis, with visual stimulus location as the predictor variable. Any participant for whom the stimulus location significantly predicted peak saccade location was not invited back for the MRI sessions. In this way, participants with stimulus-driven saccades were excluded from the study (seven older adults, three younger; see Participants subsection).

### Analysis of behavioural data

#### Auditory spatial classification task (outside the MRI scanner)

For each observer, we computed the proportion of ‘perceived right’ for each of the ten locations. These ten data points can be predicted by the psychometric function *ψ*:

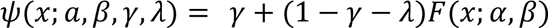

with

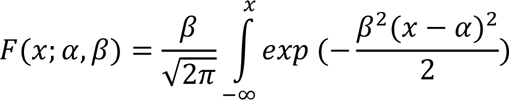

Using a Nelder-Mead optimisation algorithm, as implemented in the Palamedes toolbox (Version 1.10.3) [67] for MATLAB, we fitted a four-parameter (*α*, *β*, *λ*, *γ*) cumulative Gaussian function to these data. The parameters of this function were the mean of the distribution *α* (i.e. point of subjective equality, PSE), the slope parameter *β*, (i.e. the reciprocal of the participant’s spatial uncertainty), and the lapse parameters *λ* and *γ* (i.e. the probability of incorrectly responding right when stimuli were perceived to be on the left, and vice versa). We calculated each participant’s just-noticeable difference, a measure of spatial uncertainty, as the reciprocal of the fitted slope (JND = 1/*β*).

The point of subjective equality (PSE) and the just noticeable difference (JND) for each subject were entered into separate independent-samples *t*-tests to compare older and younger adults. Note that a JND of less than 10° was specified as an inclusion criterion (all participants met this requirement). Results of equivalent Bayesian analyses are included in the Supporting Information.

#### Spatial ventriloquist paradigm (inside and outside the MRI scanner)

For each participant, we calculated the mean auditory localisation response for each combination of auditory and visual locations. To reduce the complexity of the analyses, we pooled over the two hemifields by multiplying the average localisation responses to stimuli where the sound was in the left hemifield with -1. Likewise, we pooled the response times over the two hemifields. Hence, instead of four auditory locations, we modelled eccentricity with two levels (large ±15° versus small ±5° visual angle). Subject-specific mean auditory localisation responses were entered into a 2 (eccentricity: small, large) x 3 (sensory context: unisensory auditory, audiovisual congruent, audiovisual incongruent) x 2 (group: younger, older) mixed ANOVA with the group factor as the only between-subjects factor. Please note that for audiovisual incongruent trials, small and large eccentricity directly maps onto small and large audiovisual spatial disparity. A similar mixed ANOVA was conducted on participants’ condition-specific median reaction times (again pooled over hemifields). Results of equivalent Bayesian analyses are included in the Supporting Information.

### MRI data acquisition

A 3T Philips MRI scanner with 32-channel head coil was used to acquire both T1-weighted anatomical images (TR = 8.4 ms, TE = 3.8 ms, flip angle = 8°, FOV = 288 mm x 232 mm, image matrix = 288 x 232, 175 sagittal slices acquired in ascending direction, voxel size = 1 x 1 x 1 mm) and T2*-weighted axial echoplanar images with bold oxygenation level-dependent (BOLD) contrast (gradient echo, SENSE factor of 2, TR = 2800 ms, TE = 40 ms, flip angle = 90°, FOV = 192 mm x 192 mm, image matrix 76 x 76, 38 transversal slices acquired in ascending direction, voxel size = 2.5 x 2.5 x 2.5 mm with a 0.5 mm interslice gap).

Each participant took part in two one-hour scanning sessions, performed on separate days. In total (pooled over the two days), eleven task runs of 115 volumes each were acquired (i.e. 1265 scanning volumes in total). Each scanning session also involved a further 115- volume resting-state run, during which participants were instructed to fixate a central cross.

Four additional volumes were discarded from each scanning run prior to the analysis to allow for T_1_ equilibration effects.

### fMRI data analysis

Our fMRI analysis assessed the commonalities and differences in audiovisual spatial processing and integration between younger and older adults by combining three complementary methodological approaches. First, we used multivariate pattern decoding with support vector regression to characterise how auditory and visual information are combined into spatial representations along the dorsal visual and auditory processing hierarchies in younger and older participants. Second, we used conventional mass-univariate analyses to investigate how congruent and incongruent audiovisual stimulation influences univariate BOLD responses across the entire brain. Third, we used multivariate Bayesian decoding (MVB) to assess how the neural systems that show greater activations for older adults, as well as those that were activated in both groups, encode information about the spatial location or congruency of audiovisual stimuli.

### Preprocessing and within-subject (first-level) general linear models

MRI data were analysed in SPM12 [68]. Each participant’s functional scans were realigned/unwarped to correct for movement, slice-time corrected, and coregistered to the anatomical scan. For multivariate pattern decoding (i.e. support vector regression and multivariate Bayesian decoding), these native-space data were spatially smoothed with a Gaussian kernel of 3mm FWHM. For mass-univariate analyses and multivariate Bayesian decoding, the slice-time-corrected and realigned images were normalised into Montreal Neurological Institute (MNI) space using parameters from segmentation of the T1 structural image [69], resampled to a spatial resolution of 2 x 2 x 2 mm^3^ and spatially smoothed with a Gaussian kernel of 8 mm full-width at half-maximum.

The following processing steps were conducted separately on both native-space and MNI-transformed data. Each voxel’s time series was high-pass filtered to 1/128Hz. The fMRI experiment was modelled in an event-related fashion with regressors entered into the design matrix after convolving each event-related unit impulse (coding the stimulus onset) with a canonical hemodynamic response function and its first temporal derivative. In addition to modelling the 12 conditions in our 4 (auditory location: -15°, -5°, 5°, or 15° visual angle) x 3 (sensory context: unisensory auditory, audiovisual congruent, audiovisual incongruent) within-subject factorial design, the model included the realignment parameters as nuisance covariates to account for residual motion artifacts. For the mass-univariate analysis and the multivariate Bayesian decoding analysis, the design matrix also modelled the button response choices as a single regressor to account for motor responses. To enable more reliable estimates of the activation patterns, we did not account for observers’ response choices in the support vector regression analysis that is reported in this manuscript (sound locations and observers’ sound localisation responses were highly correlated). However, a control analysis confirmed that the fMRI decoded spatial locations did not differ across age groups when observers’ spatially specific responses were also modelled.

### Correcting BOLD response for age-related changes in vascular reactivity

To account for age-related changes in vascular reactivity, we corrected the BOLD-response amplitude (i.e. parameter estimates pertaining to the canonical hemodynamic response function) in each voxel in the MNI-normalised data based on the resting state fluctuation amplitude (RSFA or scan-to-scan signal variability)[70, 71]. Resting-state data were preprocessed exactly as the task (i.e. spatial ventriloquist) data (i.e realigned/unwarped, slice-time corrected, coregistered to the anatomical image, normalised to MNI space, resampled, and spatially smoothed with a Gaussian kernel of 8 mm FWHM). We applied additional steps to minimise the effect of motion, and other nuisance variables, on the signal. First, we applied wavelet despiking [72] and linear and quadratic detrending. The BOLD response over scans was then residualised with respect to the following regressors: white matter signal (the mean across all voxels containing white matter, according to SPM’s automated segmentation algorithm, was taken for each volume, and the time-varying signal included as a regressor); cerebrospinal fluid signal (using the same procedure as with white matter); and movement parameters (and their first derivatives). The signal was then bandpass-filtered at 0.01-0.08Hz to maximise the contribution of physiological factors to the signal fluctuation. The standard deviation of the remaining variation across scans at each voxel was calculated to create the final RSFA map (separately for each scanning day). The parameter estimates in each voxel, condition and subject were standardised by dividing by the relevant RSFA value prior to further analysis.

### Decoding audiovisual spatial estimates using support vector regression

Using multivariate pattern decoding with support vector regression (SVR), we investigated how younger and older adults combine auditory and visual signals into spatial representations along the auditory and visual processing hierarchies. The basic rationale of this analysis is as follows: We first train a model to learn the mapping from fMRI activation patterns in regions of interest to stimulus locations in the external world based solely on congruent audiovisual stimuli. We then use this learnt mapping to decode the spatial locations from activation patterns of the incongruent audiovisual signals. In putatively unisensory auditory regions, locations decoded from fMRI activation patterns for incongruent trials should therefore reflect only the sound location (irrespective of the visual location); in unisensory visual regions, decoded locations should reflect only the visual location; and in audiovisual integration regions, the decoded locations should be somewhere between the auditory and visual locations. Hence, the locations decoded from activation patterns for audiovisual incongruent stimuli provide insights into how regions combine spatial information from vision and audition.

For the multivariate decoding analysis, we extracted the parameter estimates of the canonical hemodynamic response function for each condition and run from voxels of the regions of interest (i.e. fMRI activation vectors; see definition of region of interest section below). The parameter estimates pertaining to the canonical hemodynamic response function defined the magnitude of the BOLD response to the auditory and audiovisual stimuli in each voxel. Each fMRI activation vector for the 12 conditions in our 4 (auditory location) x 3 (sensory context) factorial design was based on 10 trials within a particular run. Activation vectors were normalised to between zero and one.

For each of the five regions of interest along the visual and auditory processing hierarchies we trained a SVR model (with default parameters *C* = 1 and γ = 1/n features, as implemented in LIBSVM 3.17 [73], accessed via The Decoding Toolbox Version 3.96 [74]) to learn the mapping from the fMRI activation vectors to the external spatial locations based on the audiovisual spatially *congruent* conditions from all but one of the 11 runs. This learnt mapping from activation patterns to external spatial locations was then used to decode the spatial location from the fMRI activation patterns of the unisensory auditory, audiovisual congruent, and audiovisual incongruent conditions of the remaining run. In a leave-one-run-out cross-validation scheme, the training-test procedure was repeated for all 11 runs. The decoded spatial estimates for each condition were then averaged across runs. As in the behavioural analysis, we pooled over hemifield by multiplying the decoded spatial estimates from trials that presented auditory stimuli in the left hemifield with -1. We entered these condition-specific decoded spatial estimates (pooled over hemifields) into a 2 (eccentricity: small [±5°], large [±15°] visual angle) x 3 (sensory context: unisensory auditory, audiovisual congruent, audiovisual incongruent) x 2 (age: younger, older participants) mixed ANOVA at the second (random effects) level separately for each region of interest. For analyses and plotting, incongruent conditions were labelled based on the location of the stimulus that corresponds with the ROI’s dominant sensory modality: V1 - V3, IPS 0-2, and IPS3-4 responses were labelled based on the location of the visual stimulus; PT and A1 were labelled based on the location of the auditory stimulus. Results of equivalent Bayesian analyses are included in the Supporting Information.

### Regions of interest for SVR analysis

For the SVR analyses, all five regions of interest (ROI) were defined based on inverse-normalised group-level probabilistic maps along the auditory and visual processing streams, consistent with our previous research [5,10,13,14,51]. Left and right hemisphere maps were combined. Visual (V1 – V3) and intraparietal sulcus (IPS 0 – 2, IPS 3 – 4) ROIs were defined using retinotopic maximum probability maps [75]. Primary auditory cortex (A1) was defined based on cytoarchitectonic maximum probability maps [76]. The planum temporale (PT) was defined based on labels of the Destrieux atlas [77, 78], as implemented in Freesurfer 5.3.0 [79].

### Conventional second-level mass-univariate analysis

Using conventional mass-univariate analysis, we next characterised activations for audiovisual stimuli relative to fixation, and audiovisual spatial incongruence, across the entire brain, and compared between older and younger participants. At the first level, condition-specific effects for each participant were estimated according to the general linear model (see earlier section) and passed to a second-level ANOVA as contrasts. Inferences were made at the second level to allow for random effects analysis and population-level inferences [80].

At the random effects (i.e. group) level we tested for:

1. Effects present in both age groups for all stimuli (unisensory auditory, audiovisual congruent, and audiovisual incongruent) relative to fixation:

• (All_Older_ > Fixation_Older_) ∩ (All_Younger_ > Fixation_Younger_)
2. Age group differences in the effects of all stimuli relative to fixation:

• (All_Older_ > Fixation_Older_) > (All_Younger_ > Fixation_Younger_)
• (All_Younger_ > Fixation_Younger_) > (All_Older_ > Fixation_Older_)
3. The effect of audiovisual spatial incongruence, averaged across age groups:

• Incong > Cong
4. The interaction between audiovisual spatial incongruence and age group:

• (Incong_Older_ > Cong_Older_) > (Incong_Younger_ > Cong_Younger_)
• (Incong_Younger_ > Cong_Younger_) > (Incong_Older_ > Cong_Older_)

Unless otherwise stated, activations are reported at *p* < .05 at the voxel level, family-wise error (FWE) corrected for multiple comparisons across the entire brain.

### Multivariate Bayesian decoding

We assessed the extent to which activations identified by the mass-univariate analysis contributed to encoding of visual or auditory location, and their spatial relationship, in younger and older participants. Our key question was whether regions with greater activations for older than younger adults contribute more to encoding these task-relevant variables.

To address this question, we used multivariate Bayesian Decoding (MVB), as implemented in SPM12 [34], which estimates the set of activation patterns that best predicts a particular target variable such as visual or auditory location using hierarchical parametric empirical Bayes. Critically, because each MVB model predicts a target variable (e.g. auditory location left vs. right) based on activation patterns, we can assess the relative contributions of different ROIs to encoding a particular target variable using Bayesian model comparison. In other words, we can use standard procedures of Bayesian model comparison to assess whether activation patterns in specific regions or sets of regions are better at encoding environmental properties. In particular, MVB allows us to ask whether areas with increased BOLD responses in older than younger adults make a critical contribution to information encoding (see below).

Because the decoding of a target variable based on a large number of voxel activations (relative to a small number of scans) is an ill-posed problem, MVB imposes priors on the pattern weights at the second level of the hierarchical model. The model also includes an overall sparsity (hyper) prior accommodating the assumption that only a few patterns make a large contribution to predicting the target variable. The pattern weights (i.e. the unknown parameters defining the mapping between activation pattern and target variable) are assigned to nested sets, in which each pattern within a subset has equal variance. A greedy search algorithm iteratively optimises this nested partitioning of pattern weights to maximise the free energy as an approximation to the log model evidence. MVB estimation furnishes the log evidence for a particular model that embodies a hypothesis about the relationship between patterns of voxel activation and a target variable [34, 81]. The model evidence can then be used to compare different models using Bayesian model selection (BMS) at the group (i.e. random effects) level [82].

Specifically, we used MVB to compare representations of stimulus properties between three functionally defined ROIs:

1. Activations that are common to younger and older participants (referred to in the following as O∩Y), as specified by the conjunction (using the conjunction null [35, 36]): *(All_Older_ > Fixation_Older_) ∩ (All_Younger_ > Fixation_Younger_)*.
2. Activations that were enhanced for older relative to younger participants (referred to as [O>Y]), as specified by: *(All_Older_ > Fixation_Older_) > (All_Younger_ > Fixation_Younger_)*.
3. The union [O>Y ∪ O∩Y] of each of the above two ROIs.

These regions of interest were defined based on the respective inverse normalised statistical comparisons at the random effects group level, using a leave-one-participant-out scheme. They were constrained to include only the 1000 voxels with the greatest *t* value for the respective comparisons; the union ROI [O>Y ∪ O∩Y] was created by randomly sampling 500 voxels from each of the two component ROIs.

For each ROI we fitted four independent MVB models, predicting different target variables:

1. Visual location (VisL ≠ VisR)
2. Auditory location (AudL ≠ AudR)
3. Incongruency with 5° eccentricity (Incong5 ≠ Cong5)
4. ncongruency with 15° eccentricity (Incong15 ≠ Cong15)

Both predictor and target variables were residualised with respect to effects of no interest (i.e. all GLM covariates other than those involved in the target contrast).

The MVB analysis thus included the following steps:

First, we assessed whether information is encoded in a more sparse or distributed fashion in each region by comparing models in which patterns are individual voxels (i.e. ‘sparse’) versus clusters (i.e. smooth spatial prior). In our data the sparse model (in which the weights of individual voxels are optimised) outperformed the smooth model across all analyses (paired-sample *t-*tests of log model evidences, *p <* .001), so we will focus selectively on the results from this model class.

We also ensured that the target variables could be decoded reliably from each ROI by comparing the evidence for each ‘model of interest’ with the evidence of models in which the design matrix had been randomly phase shuffled (i.e. stimulus onset times uniformly shifted by a random amount; this was repeated 20 times, and the mean of the log model evidence was taken; see e.g. [56] for a similar approach). Using *t* tests, we compared the difference in real versus shuffled model evidences and confirmed that the real models performed significantly better for all ROIs and target variables (*p* < .05, one tailed) except Incong15 ≠ Cong15 in the O∩Y ROI, *t*(31) = 1.24, *p* = .113.

Next, and more importantly, we assessed which of the three candidate ROIs (i.e. 1. [O∩Y], the conjunction of activations in older and younger; 2. [O>Y], activation increases in older relative to younger adults; or 3. [O>Y ∪ O∩Y], the union of ROIs 1 and 2) is the best model or predictor for each of the target variables, separately for the older and younger groups, by performing Bayesian model selection at the random effects (group) level, as implemented in SPM12 [82]. We report log model evidence values, as well as the protected exceedance probability that a given model is better than any of the other candidate models beyond chance [83]. If the regions with greater activations in older (relative to younger) adults make critical contributions to encoding the task-relevant target variable, we would expect the model evidence for the union [O>Y ∪ O∩Y] to exceed that of the conjunction model [O∩Y]. Further, we formally assessed whether the frequency with which each ROI model “won” differed between age groups using a *χ*² test of association (one test per target variable). We report *p* values after Bonferroni correction for multiple (i.e. four target variables) comparisons.

Finally, we investigated whether the set of regions with greater activations for older participants (i.e. [O>Y] ROI) contributes more to the encoding of the critical target variables in older adults by comparing the difference in log model evidence for the union [O>Y ∪ O∩Y] ROI relative to the joint [O∩Y] ROI between older and younger adults in a non-parametric Mann-Whitney U tests separately for each of the four target variables (VisL ≠ VisR, AudL ≠ AudR, Incong5 ≠ Cong5, and Incong15 ≠ Cong15). We report *p* values after Bonferroni correction for multiple (i.e. four target variables) comparisons. Full output from these and the above-mentioned *χ*² tests, as well as Bayesian equivalents, are available in the Supporting Information.

## Supporting information

Supplementary Results

## Acknowledgements

The authors wish to thank Susan Francis and Stephen Mayhew for helpful discussions and support during the design of this research.

## Notes

### Competing Interest Statement

The authors have declared no competing interest.

