## Supplementary Results for "Audiovisual integration is preserved in older adults across the cortical hierarchy"

**Notes**

- All Bayesian analyses were conducted as implemented in JASP v0.16.4, using default settings unless otherwise specified
- In all cases, Bayes factors presented are $BF_{01}$ or equivalent (i.e. how likely the data are to occur under the null hypothesis, relative to a non-directional alternative hypothesis)
- Classical (Frequentist) Mixed ANOVAs are Greenhouse-Geisser corrected for sphericity violations where appropriate

### Auditory spatial classification performance for older and younger adults

***Bayesian Independent-Samples t Test***

| | $BF_{01}$ | error % |
|---|---|---|
| JND | 1.228 | 0.004 |
| PSE | 2.673 | 0.003 |

**Audiovisual integration behaviour for older and younger adults (outside the scanner)**

Results of mixed ANOVAs on mean auditory localisation responses and median reaction times in the spatial ventriloquist task during the behavioural screening session.

***Classical (Frequentist) Mixed ANOVAs***

| | df | | | | |
|---|---|---|---|---|---|
| | effect | error | *F* | *p* | $\eta_p^2$ |
| *Mean localisation responses* | | | | | |
| Eccentricity | 1 | 30 | 250.132 | < .001 | .893 |
| Eccentricity x Age | 1 | 30 | 2.528 | .122 | .078 |
| Congruence | 1.146 | 34.377 | 47.559 | < .001 | .613 |
| Congruence x Age | 1.146 | 34.377 | 1.140 | .302 | .037 |
| Eccentricity x Sensory context | 1.268 | 38.046 | 1.295 | .272 | .041 |
| Eccentricity x Sensory context x Age | 1.268 | 38.046 | 1.707 | .201 | .054 |
| Age | 1 | 30 | 3.556 | .069 | .106 |
| *Median reaction times* | | | | | |
| Eccentricity | 1 | 30 | 22.588 | < .001 | .430 |
| Eccentricity x Age | 1 | 30 | 4.253 | .048 | .124 |
| Sensory context | 2 | 60 | 30.845 | < .001 | .507 |
| Sensory context x Age | 2 | 60 | 1.336 | .271 | .043 |
| Eccentricity x Sensory context | 2 | 60 | 7.164 | .002 | .193 |
| Eccentricity x Sensory context x Age | 2 | 60 | 2.012 | .143 | .063 |
| Age | 1 | 30 | 16.358 | < .001 | .353 |

***Bayesian Mixed ANOVAs***

| Effects | P(incl) | P(excl) | P(incl\|data) | P(excl\|data) | $BF_{excl}$ |
|---|---|---|---|---|---|
| *Mean localisation responses* | | | | | |
| Eccentricity | 0.737 | 0.263 | 1.000 | $9.925 \times 10^{-14}$ | $2.779 \times 10^{-13}$ |
| Age | 0.737 | 0.263 | 0.721 | 0.279 | 1.083 |
| Eccentricity x Age | 0.316 | 0.684 | 0.262 | 0.738 | 1.297 |
| Sensory context | 0.737 | 0.263 | 1.000 | $1.999 \times 10^{-11}$ | $5.597 \times 10^{-11}$ |
| Eccentricity x Sensory context | 0.316 | 0.684 | 0.254 | 0.746 | 1.353 |
| Age x Sensory context | 0.316 | 0.684 | 0.219 | 0.781 | 1.647 |
| Eccentricity x Age x Sensory context | 0.053 | 0.947 | 0.015 | 0.985 | 3.636 |
| *Median reaction times* | | | | | |
| Eccentricity | 0.737 | 0.263 | 1.000 | $8.244 \times 10^{-5}$ | $2.309 \times 10^{-4}$ |
| Sensory context | 0.737 | 0.263 | 1.000 | $1.471 \times 10^{-9}$ | $4.119 \times 10^{-9}$ |
| Eccentricity x Sensory context | 0.316 | 0.684 | 0.952 | 0.048 | 0.023 |
| Age | 0.737 | 0.263 | 0.995 | 0.005 | 0.013 |
| Eccentricity x Age | 0.316 | 0.684 | 0.655 | 0.345 | 0.243 |
| Sensory context x Age | 0.316 | 0.684 | 0.380 | 0.620 | 0.752 |
| Eccentricity x Sensory context x Age | 0.053 | 0.947 | 0.114 | 0.886 | 0.433 |

**Audiovisual integration behaviour for older and younger adults (inside the scanner)**

***Bayesian Mixed ANOVAs***

| Effects | P(incl) | P(excl) | P(incl\|data) | P(excl\|data) | $BF_{excl}$ |
|---|---|---|---|---|---|
| *Mean localisation responses* | | | | | |
| Eccentricity | 0.737 | 0.263 | 1.000 | $7.305\times10^{-14}$ | $2.045\times10^{-13}$ |
| Sensory context | 0.737 | 0.263 | 1.000 | $3.407\times10^{-12}$ | $9.540\times10^{-12}$ |
| Eccentricity x Sensory context | 0.316 | 0.684 | 0.444 | 0.556 | 0.578 |
| Age | 0.737 | 0.263 | 0.732 | 0.268 | 1.025 |
| Eccentricity x Age | 0.316 | 0.684 | 0.444 | 0.556 | 0.578 |
| Sensory context x Age | 0.316 | 0.684 | 0.338 | 0.662 | 0.903 |
| Eccentricity x Sensory context x Age | 0.053 | 0.947 | 0.197 | 0.803 | 0.227 |
| *Median reaction times* | | | | | |
| Eccentricity | 0.737 | 0.263 | 0.932 | 0.068 | 0.205 |
| Sensory context | 0.737 | 0.263 | 1.000 | $1.902\times10^{-9}$ | $5.326\times10^{-9}$ |
| Eccentricity x Sensory context | 0.316 | 0.684 | 0.837 | 0.163 | 0.090 |
| Age | 0.737 | 0.263 | 0.968 | 0.032 | 0.093 |
| Eccentricity x Age | 0.316 | 0.684 | 0.309 | 0.691 | 1.031 |
| Sensory context x Age | 0.316 | 0.684 | 0.563 | 0.437 | 0.358 |
| Eccentricity x Sensory context x Age | 0.053 | 0.947 | 0.066 | 0.934 | 0.780 |

**Decoding audiovisual spatial estimates using support vector regression (analyses <u>including</u> unisensory auditory condition)**

***Bayesian Mixed ANOVAs***

| Effects | P(incl) | P(excl) | P(incl\|data) | P(excl\|data) | $BF_{excl}$ |
|---|---|---|---|---|---|
| *V1- 3* | | | | | |
| Eccentricity | 0.737 | 0.263 | 1.000 | $1.887\times10^{-15}$ | $5.285\times10^{-15}$ |
| Sensory context | 0.737 | 0.263 | 1.000 | $1.443\times10^{-15}$ | $4.041\times10^{-15}$ |
| Eccentricity x Sensory context | 0.316 | 0.684 | 1.000 | $2.286\times10^{-7}$ | $1.055\times10^{-7}$ |
| Age | 0.737 | 0.263 | 0.800 | 0.200 | 0.698 |
| Eccentricity x Age | 0.316 | 0.684 | 0.396 | 0.604 | 0.704 |
| Sensory context x Age | 0.316 | 0.684 | 0.708 | 0.292 | 0.190 |
| Eccentricity x Sensory context x Age | 0.053 | 0.947 | 0.242 | 0.758 | 0.174 |
| *IPS 0 -2* | | | | | |
| Eccentricity | 0.737 | 0.263 | 1.000 | $5.820\times10^{-8}$ | $1.630\times10^{-7}$ |
| Sensory context | 0.737 | 0.263 | 1.000 | 0.000 | 0.000 |
| Age | 0.737 | 0.263 | 0.497 | 0.503 | 2.833 |
| Eccentricity x Sensory context | 0.316 | 0.684 | 0.999 | 0.001 | $6.824\times10^{-4}$ |
| Eccentricity x Age | 0.316 | 0.684 | 0.180 | 0.820 | 2.103 |
| Sensory context x Age | 0.316 | 0.684 | 0.079 | 0.921 | 5.358 |
| Eccentricity x Sensory context x Age | 0.053 | 0.947 | 0.009 | 0.991 | 5.998 |
| *IPS 3 - 4* | | | | | |
| Eccentricity | 0.737 | 0.263 | 0.998 | 0.002 | 0.007 |
| Sensory context | 0.737 | 0.263 | 1.000 | $2.514\times10^{-6}$ | $7.039\times10^{-6}$ |
| Age | 0.737 | 0.263 | 0.344 | 0.656 | 5.349 |
| Eccentricity x Sensory context | 0.316 | 0.684 | 0.996 | 0.004 | 0.002 |
| Eccentricity x Age | 0.316 | 0.684 | 0.106 | 0.894 | 3.881 |
| Sensory context x Age | 0.316 | 0.684 | 0.078 | 0.922 | 5.485 |
| Eccentricity x Sensory context x Age | 0.053 | 0.947 | 0.010 | 0.990 | 5.695 |
| *PT* | | | | | |
| Eccentricity | 0.737 | 0.263 | 1.000 | $1.783\times10^{-4}$ | $4.994\times10^{-4}$ |
| Sensory context | 0.737 | 0.263 | 0.995 | 0.005 | 0.013 |
| Age | 0.737 | 0.263 | 0.347 | 0.653 | 5.269 |
| Eccentricity x Sensory context | 0.316 | 0.684 | 0.426 | 0.574 | 0.621 |
| Eccentricity x Age | 0.316 | 0.684 | 0.083 | 0.917 | 5.117 |
| Sensory context x Age | 0.316 | 0.684 | 0.082 | 0.918 | 5.185 |
| Eccentricity x Sensory context x Age | 0.053 | 0.947 | 0.002 | 0.998 | 32.444 |
| *A1* | | | | | |
| Eccentricity | 0.737 | 0.263 | 0.998 | 0.002 | 0.005 |
| Sensory context | 0.737 | 0.263 | 0.735 | 0.265 | 1.010 |
| Age | 0.737 | 0.263 | 0.320 | 0.680 | 5.955 |
| Eccentricity x Sensory context | 0.316 | 0.684 | 0.066 | 0.934 | 6.480 |
| Eccentricity x Age | 0.316 | 0.684 | 0.082 | 0.918 | 5.193 |
| Sensory context x Age | 0.316 | 0.684 | 0.047 | 0.953 | 9.299 |
| Eccentricity x Sensory context x Age | 0.053 | 0.947 | $1.728\times10^{-4}$ | 1.000 | 321.399 |

**Decoding audiovisual spatial estimates using support vector regression (analyses excluding unisensory auditory condition)**

***Classical (Frequentist) Mixed ANOVAs***

| | *df* | | | | |
|---|---|---|---|---|---|
| | effect | error | *F* | *p* | $\eta_p^2$ |
| *V1 - 3* | | | | | |
| Eccentricity | 1 | 30 | 88.921 | < .001 | .748 |
| Eccentricity x Age | 1 | 30 | 2.353 | .135 | .073 |
| Congruence | 1 | 30 | 23.857 | < .001 | .443 |
| Congruence x Age | 1 | 30 | 3.687 | .064 | .109 |
| Eccentricity x Congruence | 1 | 30 | 7.154 | .012 | .193 |
| Eccentricity x Congruence x Age | 1 | 30 | 0.301 | .587 | .010 |
| Age | 1 | 30 | 1.927 | .175 | .060 |
| *IPS 0 - 2* | | | | | |
| Eccentricity | 1 | 30 | 66.462 | < .001 | .689 |
| Eccentricity x Age | 1 | 30 | 3.467 | .072 | .104 |
| Congruence | 1 | 30 | 5.407 | .027 | .153 |
| Congruence x Age | 1 | 30 | 0.213 | .648 | .007 |
| Eccentricity x Congruence | 1 | 30 | 8.860 | .006 | .228 |
| Eccentricity x Congruence x Age | 1 | 30 | 1.244 | .274 | .040 |
| Age | 1 | 30 | 0.862 | .361 | .028 |
| *IPS 3 - 4* | | | | | |
| Eccentricity | 1 | 30 | 1.314 | .261 | .042 |
| Eccentricity x Age | 1 | 30 | 0.278 | .602 | .009 |
| Congruence | 1 | 30 | 21.997 | < .001 | .423 |
| Congruence x Age | 1 | 30 | 0.142 | .709 | .005 |
| Eccentricity x Congruence | 1 | 30 | 12.422 | < .001 | .293 |
| Eccentricity x Congruence x Age | 1 | 30 | 1.254 | .272 | .040 |
| Age | 1 | 30 | 0.117 | .734 | .004 |
| *PT* | | | | | |
| Eccentricity | 1 | 30 | 36.481 | < .001 | .549 |
| Eccentricity x Age | 1 | 30 | 0.782 | .782 | .003 |
| Congruence | 1 | 30 | 23.168 | < .001 | .436 |
| Congruence x Age | 1 | 30 | 0.050 | .825 | .002 |
| Eccentricity x Congruence | 1 | 30 | 1.332 | .258 | .043 |
| Eccentricity x Congruence x Age | 1 | 30 | 0.477 | .495 | .016 |
| Age | 1 | 30 | 0.023 | .880 | .001 |
| *A1* | | | | | |
| Eccentricity | 1 | 30 | 17.913 | < .001 | .374 |
| Eccentricity x Age | 1 | 30 | 0.169 | .684 | .006 |
| Congruence | 1 | 30 | 5.346 | .028 | .151 |
| Congruence x Age | 1 | 30 | 1.056 | .312 | .034 |
| Eccentricity x Congruence | 1 | 30 | 0.001 | .969 | < .001 |
| Eccentricity x Congruence x Age | 1 | 30 | 0.181 | .674 | .006 |
| Age | 1 | 30 | 0.093 | .763 | .003 |

***Bayesian Mixed ANOVAs***

| Effects | P(incl) | P(excl) | P(incl\|data) | P(excl\|data) | $BF_{excl}$ |
|---|---|---|---|---|---|
| *V1 - 3* | | | | | |
| Eccentricity | 0.737 | 0.263 | 1.000 | $1.751\times10^{-9}$ | $4.903\times10^{-9}$ |
| Sensory context | 0.737 | 0.263 | 1.000 | $4.033\times10^{-4}$ | 0.001 |
| Age | 0.737 | 0.263 | 0.785 | 0.215 | 0.768 |
| Eccentricity x Sensory context | 0.316 | 0.684 | 0.843 | 0.157 | 0.086 |
| Eccentricity x Age | 0.316 | 0.684 | 0.404 | 0.596 | 0.681 |
| Sensory context x Age | 0.316 | 0.684 | 0.425 | 0.575 | 0.623 |
| Eccentricity x Sensory context x Age | 0.053 | 0.947 | 0.057 | 0.943 | 0.920 |
| *IPS 0 – 2* | | | | | |
| Eccentricity | 0.737 | 0.263 | 1.000 | $2.442\times10^{-7}$ | $6.836\times10^{-7}$ |
| Sensory context | 0.737 | 0.263 | 0.987 | 0.013 | 0.038 |
| Age | 0.737 | 0.263 | 0.523 | 0.477 | 2.553 |
| Eccentricity x Sensory context | 0.316 | 0.684 | 0.954 | 0.046 | 0.022 |
| Eccentricity x Age | 0.316 | 0.684 | 0.247 | 0.753 | 1.405 |
| Sensory context x Age | 0.316 | 0.684 | 0.168 | 0.832 | 2.281 |
| Eccentricity x Sensory context x Age | 0.053 | 0.947 | 0.037 | 0.963 | 1.455 |
| *IPS 3 – 4* | | | | | |
| Eccentricity | 0.737 | 0.263 | 0.997 | 0.003 | 0.009 |
| Sensory context | 0.737 | 0.263 | 1.000 | $6.587\times10^{-5}$ | $1.845\times10^{-4}$ |
| Age | 0.737 | 0.263 | 0.366 | 0.634 | 4.843 |
| Eccentricity x Sensory context | 0.316 | 0.684 | 0.996 | 0.004 | 0.002 |
| Eccentricity x Age | 0.316 | 0.684 | 0.110 | 0.890 | 3.727 |
| Sensory context x Age | 0.316 | 0.684 | 0.102 | 0.898 | 4.058 |
| Eccentricity x Sensory context x Age | 0.053 | 0.947 | 0.014 | 0.986 | 3.815 |
| *PT* | | | | | |
| Eccentricity | 0.737 | 0.263 | 1.000 | $8.035\times10^{-5}$ | $2.250\times10^{-4}$ |
| Sensory context | 0.737 | 0.263 | 0.997 | 0.003 | 0.007 |
| Age | 0.737 | 0.263 | 0.358 | 0.642 | 5.011 |
| Eccentricity x Sensory context | 0.316 | 0.684 | 0.372 | 0.628 | 0.779 |
| Eccentricity x Age | 0.316 | 0.684 | 0.090 | 0.910 | 4.687 |
| Sensory context x Age | 0.316 | 0.684 | 0.086 | 0.914 | 4.908 |
| Eccentricity x Sensory context x Age | 0.053 | 0.947 | 0.003 | 0.997 | 16.790 |
| *A1* | | | | | |
| Eccentricity | 0.737 | 0.263 | 0.996 | 0.004 | 0.012 |
| Sensory context | 0.737 | 0.263 | 0.763 | 0.237 | 0.872 |
| Age | 0.737 | 0.263 | 0.375 | 0.625 | 4.664 |
| Eccentricity x Sensory context | 0.316 | 0.684 | 0.161 | 0.839 | 2.401 |
| Eccentricity x Age | 0.316 | 0.684 | 0.108 | 0.892 | 3.818 |
| Sensory context x Age | 0.316 | 0.684 | 0.111 | 0.889 | 3.710 |
| Eccentricity x Sensory context x Age | 0.053 | 0.947 | 0.002 | 0.998 | 25.767 |

**Multivariate Bayesian decoding**

***Mann-Whitney* U *Tests of Age Differences in Model Evidence "Boost" for [O>Y ∪ O∩Y] over [O∩Y]***

| Target variable | W | p |
|---|---|---|
| VisL ≠ VisR | 116.000 | 0.669 |
| AudL ≠ AudR | 126.000 | 0.956 |
| Incong5 ≠ Cong5 | 139.000 | 0.696 |
| Incong15 ≠ Cong15 | 69.000 | 0.026 |

***Bayesian Mann-Whitney* U *Tests of Age Differences in Model Evidence "Boost" for [O>Y ∪ O∩Y] over [O∩Y]***

| Target variable | $BF_{01}$ | W | Rhat |
|---|---|---|---|
| VisL ≠ VisR | 2.415 | 116.000 | 1.000 |
| AudL ≠ AudR | 2.866 | 126.000 | 1.000 |
| Incong5 ≠ Cong5 | 2.568 | 139.000 | 1.000 |
| Incong15 ≠ Cong15 | 0.616 | 69.000 | 1.000 |

*Note.* Result based on data augmentation algorithm with 5 chains of 10000 iterations.

***$\chi^2$ Tests of Association on Age Differences in the Frequency with Which Each MVB Model "Won" – Classical and Bayesian ($BF_{01}$ Independent Multinomial)***

| Target variable | $\chi^2$ | N | df | p | $BF_{01}$ |
|---|---|---|---|---|---|
| VisL ≠ VisR | 1.091 | 32 | 2 | 0.580 | 5.926 |
| AudL ≠ AudR | 2.159 | 32 | 2 | 0.340 | 3.085 |
| Incong5 ≠ Cong5 | 0.821 | 32 | 2 | 0.365 | 2.111 |
| Incong15 ≠ Cong15 | 0.582 | 32 | 2 | 0.446 | 1.979 |
